# No signal of a super-archaic origin of the Denisovan AMBN gene, a comment on the protein affinity of *Homo erectus* and Denisovan enamel proteins

**DOI:** 10.64898/2026.08.11.744165

**Authors:** Ioannis Patramanis, Frido Welker, Laurits Skov

## Abstract

*Homo erectus* is a species that occupies a central role in the study of human evolution. To investigate its taxonomic identity, Fu et al. 2026 extracted and sequenced enamel proteins from 6 fossils assigned to *Homo erectus*, originating from 3 localities in China and dated to around 400 thousand years ago. Unexpectedly, all samples possess an amino acid variant on the enamel protein ameloblastin (AMBN) which is uniquely shared with more recent Denisovan fossils and a subset of present day humans who are known to carry introgressed Denisovan ancestry. The samples also showcase an additional, novel variant on the same enamel protein. To explain this result, Fu et al. 2026 propose a model where the sampled *Homo erectus* population (or its recent ancestors) interbred with later arriving Denisovans, introducing one of the two AMBN variants into the Denisovan population. While this model fits our overall understanding of the interactions between these archaic populations, it rests on the assumption that the Denisovan AMBN variant has an archaic, *Homo erectus-*like, source. Here we show that the AMBN gene of late Denisovans has no signal of introgression from a ‘super-archaic’ source, making the suggested model unlikely for this gene. We also show that the AMBN gene of Denisova 25, an earlier Denisova, does show a signal of introgression, but one that better matches Neanderthals and modern humans, rather than a super-archaic source. We propose a number of alternative models that could help explain the observed affinity between the sampled *H. erectus* and Denisovans without requiring the introgression of AMBN from an archaic source into Denisovans. We discuss their strengths and weaknesses and how new data could help resolve them.

## Introduction

The taxonomy of the hominin populations occupying East Asia during the Middle and Late Pleistocene is a complicated and debated topic^1^. Studies based on either morphology^2–5^ or genetics^6,7^ support the presence of multiple distinct lineages during this period, although disagreement persists on the number and the taxonomic grouping of these archaic populations. Chief among these taxonomic units is *Homo erectus*, a species initially defined based on fossils from the island of Java, but expanded to encompass samples as far as Africa, and a broad timespan of around 2 million years^8,9^. This broad definition includes multiple fossils from Asia, which are sometimes grouped together as ‘late surviving’, ‘East’^10^ or ‘Classic’^1^ *Homo erectus*. A different group of hominins, initially identified using ancient DNA from a fossil in the Denisova cave in Siberia and referred to as the “Denisovans”, have also occupied East Asia in the late Pleistocene^11^. Dental and skeletal elements identified in Tibet^12,13^, Laos ^14^, Northern China^15^ and the coast of Taiwan^16^ have been assigned to the Denisovan lineage based on morphology, ancient DNA or ancient protein analyses. Genetic studies have also shown that these Denisovan groups interbred with modern humans in the last 50.000 years^17–19^. In addition, the Denisovan genome itself contains evidence of introgression from some other, currently unidentified, hominin group^20^. This older admixture is usually referred to as “super-archaic” due to the estimated high level of divergence of the unknown hominin^21^. Given its archaic morphology, as well as the geographical and chronological overlap, *Homo erectus* fits well as the source of this older introgression.

To further illuminate our understanding of this species, Fu et al. 2026 employed the use of liquid chromatography–tandem mass spectrometry (LC–MS/MS) to sequence the enamel proteins of 6 fossils assigned to this lineage^22^. The enamel sequences of all 6 individuals showed a unique mutation on the protein of AMBN, not found in any other present or archaic population so far. This mutation changes the 253rd amino acid of the AMBN protein sequence from A>G, and can be caused by a mutation at chromosome 4:71,469,598 (hg19 coordinates). Perhaps even more intriguing was the fact that all 6 *Homo erectus* samples possess a second amino acid mutation that is, so far, found only in Denisovans (genomes or proteomes) and present day humans with significant Denisovan admixture. This second mutation changes the 273rd amino acid of the AMBN protein sequence from M>V, and is caused by a mutation at chromosome 4:71,471,920 (hg19 coordinates). In present day humans, this corresponds to the rs564905233 single nucleotide polymorphism (SNP). To explain this rather unexpected result, Fu et al. proposed a model where the rs564905233 mutation has originated in the *Homo erectus* population and then introgressed into Denisovans, later introgressing from Denisovans to present day humans in Asia (see Fig. 1A). Given the super-archaic component found in the high coverage genomes of the Denisovans from Denisova Cave^23,24^, the team further conducted an analysis to investigate whether the rs564905233 mutation in the Denisovan genome stems from a super-archaic source.

**Fig. 1.**
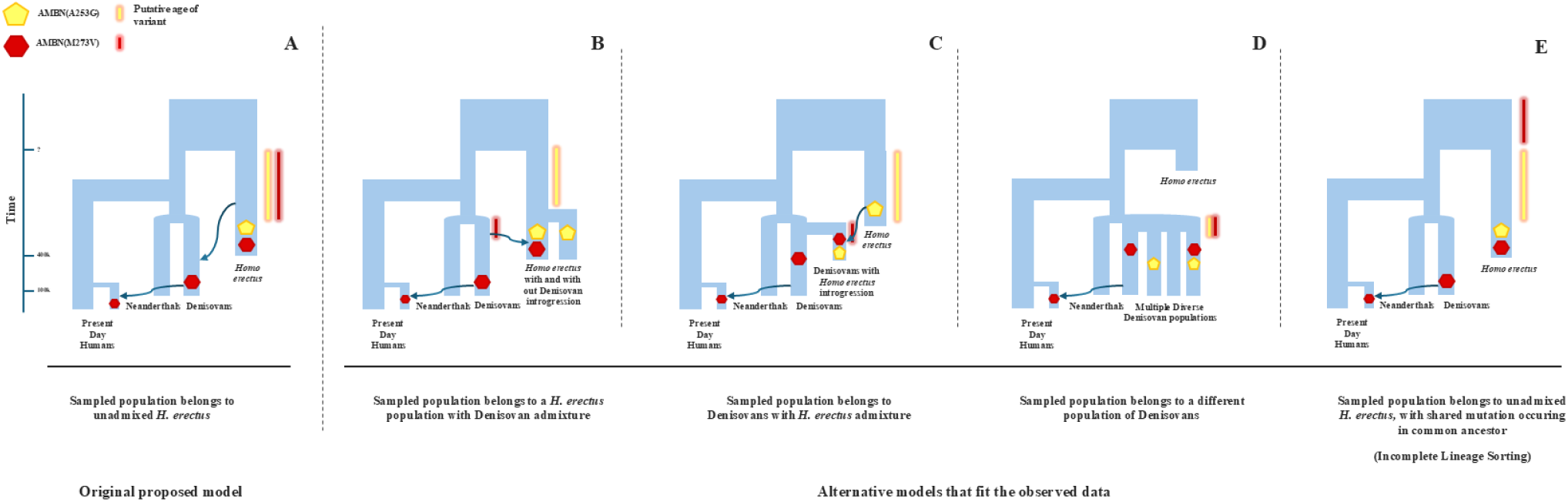
Evolutionary models comparison. Possible evolutionary models that can explain the observation of AMBN(M273V) / rs564905233 in both Denisovans, present day humans and the sequenced *Homo erectus* samples. The two coloured symbols correspond to the observation of the two mutations at a specific timepoint: AMBN(M273V) - yellow and AMBN(A253G) - red. The two coloured bars correspond to the theoretical relative timepoint and lineage, where each of the two mutations appeared. Admixture between lineages transferring AMBN are depicted with an arrow, but introgression events that are not related to AMBN are not shown.

In contrast to the rest of the genome, regions of DNA that originate from an introgression event of a deeply divergent lineage should showcase an increased amount of derived alleles, when compared with modern humans and Neanderthals. This increased amount of derived alleles would result in Denisovan DNA segments that appear more divergent to present day humans than their homologous Neanderthal segments do. For the rest of the genome, which bears no super-archaic ancestry, Denisovan and Neanderthal segments would appear roughly equidistant to present day humans. To test this, Fu et al. 2026 employed a sliding 20 kb window analysis of a 276 kb region of chromosome 4 surrounding the rs564905233 allele. They employed this analysis to study the high coverage genome of Denisova 3, which is homozygous for rs564905233. For each sliding window they calculated the difference in the genetic distance between present day Africans to Neanderthals and the genetic distance between present day Africans to Denisovans (Denisova 3). They found that rs564905233 falls within a window where this difference between the two distances is statistically significant, supported by a low p-value. They also demonstrated that the same missense SNP is also found in a heterozygous state in two older Denisovan samples, Denisova 25^24^ and the Harbin cranium^22^ (both samples date to around 150.000 to 300.000 years ago). They thus infer that this SNP (including the AMBN gene and the 20kb region surrounding it) is most likely originating from a super-archaic introgression event into the ancestor of all of these Denisovan samples. Given the protein variant resulting from this missense SNP is present in all 6 of the sequenced *Homo erectus* samples, a morphologically super-archaic lineage, they present the model of Fig. 1A as the best explanation for their results.

This model makes sense given that, firstly, the rs564905233 SNP appears to be introgressed from a super-archaic origin, and secondly, the 6 *Homo erectus* samples have a super-archaic profile, from a morphological point of view, and also have the protein variant that is the result of rs564905233.

However, there are a couple of considerations with the methodology that assigned rs564905233 to a super-archaic origin. The difference between present day Africans and Neanderthals and present day Africans and Denisovans in 20 kb windows, varies greatly across the genome. It might be the case that, in the 20 kb window surrounding the rs564905233 SNP, the similarity between present day Africans to Neanderthals is significantly different than that of present day Africans to Denisovans. Yet, while this signal appears significant in this particular 20 kb region, it is unclear how often a region of the same length (20 kb) would show such a signal. As Fu et al. already demonstrated, in the 276 kb length region of chromosome 4 where this metric was employed, one can already observe multiple significant peaks (see Supplementary material of Fu et al. 2026^22^). Furthermore, the observed difference in distance between present day Africans and Neanderthals and present day Africans and Denisovans, could be due to either incomplete lineage sorting, early modern human introgression into the Neanderthal genome, or the super-archaic introgression event into Denisovans. A more principled approach to detect super-archaic segments, would be to calculate the number of derived alleles that differ between Denisovas and Neanderthals to a present day African outgroup. This analysis should be carried out on a genome wide level, for each archaic individual, observing how the AMBN gene fits into the overall distribution of the genome.

Alternative approaches to detect potentially introgressed regions of the genome with super-archaic ancestry exist. For example, previous studies have identified regions or loci of introgression by generating phylogenetic trees of the candidate introgressed regions^25^. In the context of AMBN, if rs564905233 is the result of a super-archaic introgression event in the Denisovan genome, then one would expect the phylogenetic tree of this gene to show the Denisovan haplotypes, containing the rs564905233 SNP, as an outgroup to the haplotypes of modern humans and Neanderthals. Denisova 25, heterozygous for this SNP, should contain one super-archaic haplotype clustering with Denisova 3 and one that does not.

## Methodology

To test whether the genetic region surrounding AMBN shows signs of super-archaic ancestry, we calculated the density of derived variants present in two high coverage Denisovan genomes^23,24^ and 3 high coverage Neanderthal genomes^20,26,27^. We calculated the density in non-overlapping windows with a length of 20,000 base pairs. We counted variants that were derived with respect to the inferred ancestral allele (ftp://ftp.ensembl.org/pub/release-74/fasta/ancestral_alleles/hg19_ancestral.tar.bz2) and not present in 292 Sub-Saharan individuals from Esan, Yoruba and Menda populations from the 1000 genomes project^28^. Due to the lack of phasing we randomly sampled an allele from each high coverage archaic genome. We calculate the number of derived variants divided by the number of called bases in the 20kb window using the “joined manifesto filter” (which can be downloaded from https://cdna.eva.mpg.de/neandertal/). We then calculated empirical p-values for the derived variant density for the 20 kb region containing AMBN (chr4:71,460,000-71,480,000) and a comparative region, which has been proposed to be due to super-archaic introgression and contains the gene KRTAP10-10 (chr21:46,040,000-46,060,000)^24^. We plotted the distribution of counts for each archaic genome as a histogram (Fig. 2).

**Fig. 2.**
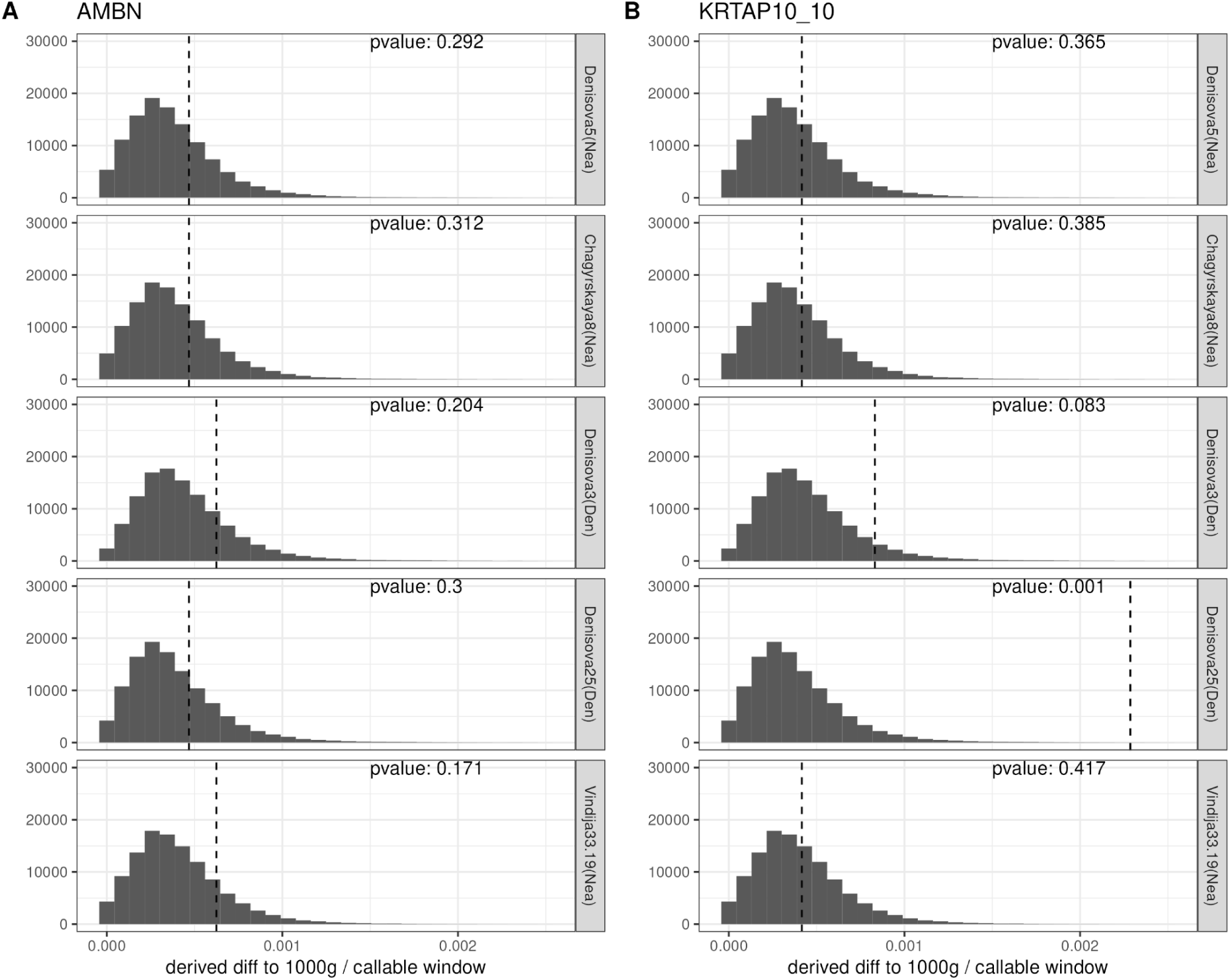
Non overlapping windows analysis. Genome wide distributions of derived alleles per 20kb, non-overlapping windows for five high coverage archaic individuals. For each individual the placement of the window containing AMBN (A) and KRTAP-10_10 (B) within the distribution is shown with a dotted vertical line.

To assess whether the Denisovan AMBN haplotype containing rs564905233 showcases the topology of a super-archaic segment, we performed a local phylogenetic analysis of the AMBN haplotypes from recent hominin groups for which high quality DNA data available. We first generated a FASTA sequence dataset of AMBN haplotypes from Neanderthals, Denisovans and present day humans. We extracted the haplotypes of present day humans from phased data of the 1000 genomes (1KG)^29^, creating two separate haplotypes for each individual. We also included 3 high coverage Neanderthal^20,26,27^ and 2 high coverage Denisovan^23,24^ genomes. For the 5 archaic individuals, which lacked phasing, we created two pseudohaplotypes per individual by assigning heterozygous sites on either pseudohaplotype. The differences between each pseudohaplotype are minimal for 4 out 5 of the archaic individuals, due to their high homozygosity. This is not the case for Denisova 25, which shows a higher heterozygosity for AMBN than any other high coverage archaic individual. Specifically, the published Denisova 25 data show a heterozygous state for 19 sites within AMBN, of which 17 have a different fixed state between the other 4 archaics and present day Africans (Extended Data Table 1). To investigate this, we created two pseudohaplotypes, assuming all archaic-like mutations to be on one haplotype and all modern-human-like mutations to be on the other. We repeated our phylogenetic analysis with and without the pseudohaplotypes of Denisova 25. We also included the reference sequence of *Pan troglodytes* from Ensembl^30^ to be used as an outgroup. After aligning the FASTA dataset using MAFFT^31^, we generated a phylogenetic tree of the AMBN gene using IQTree2^32^, rooted it using the *Pan troglodytes* AMBN sequence and plotted the results (Fig. 3).

**Fig. 3.**
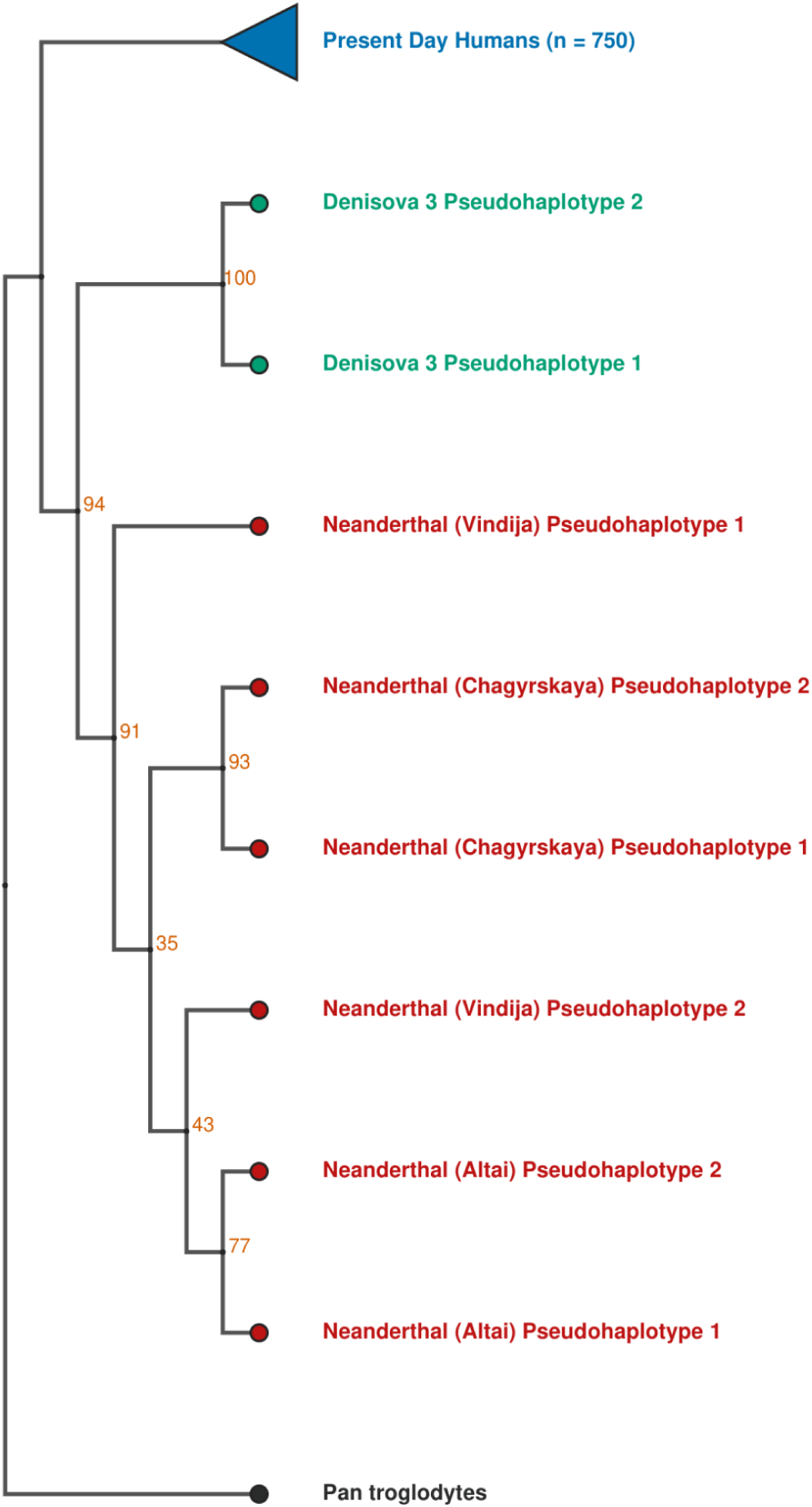
Phylogenetic tree of AMBN Haplotypes. Maximum likelihood phylogenetic tree of the AMBN haplotypes for Neanderthals, Denisova 3 and present day humans. The tree is rooted using *Pan paniscus*. Bootstrap support values of each internal node are shown in orange.

We further investigated the high levels of heterozygosity of Denisova 25 in the region around AMBN. We used a sliding window approach and calculated the genetic distance of Denisova 25 from each of the three groups (Modern humans, Neanderthals, Denisovans), as well as the number of heterozygous sites within the window. For modern humans we used the frequency of each SNP in present day humans from Africa in the Gnomad dataset^33^, for Neanderthals, the frequency in the 3 high coverage genomes^20,26,27^ and for Denisovans, the allele count in Denisova 3^23^. For each group we report the absolute distance by subtracting the allele count of Denisova 25 from each of these 3 numbers for each window. We also report the number of heterozygous sites called in each window for Denisova 25. We used a window of 20kb (roughly matching the length of the AMBN gene), sliding by 2kb, for a total region of 1Mb centered on AMBN.

## Results

The divergence between present day human individuals and Denisova 3, at the 20 kb region containing the AMBN gene, is around 6 mutations per 10,000 base pairs, placing it in the top 20% percentile of the genome wide distribution (Fig. 2A). Previous studies suggest at most 10% of super archaic introgression into Denisovans^20^. As a comparison for this region, the divergence from the Croatian Neanderthal (Vindija33.19) to present day humans is also around 6 mutations per 10,000 base pairs and there is currently no evidence of super archaic introgression into Vindija33.19. These observations do not support a scenario where the AMBN gene has a super archaic origin in Denisova 3. This is in contrast to the divergence at the KRTAP10-10 gene in Denisova 25, which has also been proposed to be due to super archaic introgression^24^. There, the divergence is around 22 mutations per 10,000 base pairs, placing it at the top 0.1% of the genomewide distribution (Fig. 2B). For Vindija33.19, for the same region, the number of mutations per 10,000 base pairs is around 4.

Our first iteration of the phylogenetic analysis of AMBN included the haplotypes of modern humans, the pseudohaplotypes of 3 Neanderthals and the pseudohaplotypes of Denisova 3, covering around 15,000 basepairs. The resulting phylogenetic tree of AMBN, mirrors that of the population-level relations between the 3 groups (Fig. 3). All three groups of Neanderthals, Denisovans and present day humans cluster in monophyletic clades with no signs of incomplete lineage sorting or admixture. Importantly, the Neanderthal and Denisova clades show a more recent common ancestor between them than either one has with present day humans. This result is contrary to what is expected of a super-archaic introgressed region, which should show the Denisova 3 AMBN haplotypes as more divergent to the clade of Neanderthals and modern humans.

When we included Denisova 25 in our phylogenetic analysis, the results changed. While the rest of the tree remains the same, the two Denisova 25 pseudohaplotypes cluster neither with Denisova 3, nor with each other. The archaic-like pseudohaplotype of Denisova 25 clusters with the 3 high coverage Neanderthals, while the modern-human-like pseudohaplotype clusters with modern humans, albeit both with low bootstrap support (Extended Data Fig. 1). When investigating this region further, we observed that this pattern of heterozygosity and similarity to modern humans and Neanderthals, and dissimilarity to Denisova 3, extends further upstream of AMBN, for a region around 100 kb (Chr4:71,372,179-71,470,419), but not downstream (see Extended Data Fig. 2). Using the length of this segment (100 kb), the recombination rate of the region (5.42e-08) and a drift time of 6,896 generations (assuming 29 years per generation, a split time of 750.000 years ago from the human lineage, and the 200.000 years date of the Denisova 25 sample) we calculated the probability that this pseudohaplotype is matching modern humans due to incomplete lineage sorting to be very low (p-value = 5.85e-17) ^34^. Other small regions further upstream of AMBN also showcase this pattern of heterozygosity matching modern humans and Neanderthals more than Denisova 3. Downstream of AMBN, the region appears archaic, but with Neanderthals and Denisova 3 having roughly equal distance to Denisova 25. In fact, the gene of AMBN seems to be located at the very tail end of this transition (Extended Data Fig. 2, Extended Data Table 1).

The results of our combined analyses do not provide strong evidence for a super-archaic origin of the rs564905233 SNP of AMBN, in either high coverage Denisovan genome. As a result, we revisit the hypothesis of an initial introgression of the rs564905233 SNP from *Homo erectus* into Denisovans, and suggest a few alternative models that could better explain the data at hand (Fig. 1B-1E).

## Discussion

Fu et al. 2026 present enamel proteomic data from 6 *Homo erectus* samples from China dating to around 400,000 years ago. The recovery of these protein sequences is groundbreaking, revealing unexpected results and showing for the first time a genetic affinity that connects these *Homo erectus* samples to the genomes of the Denisovans, and even present day humans. The authors propose a model that elegantly explains this affinity, pairing the previously identified ghost, super-archaic, component found in the Denisovan genomes to the fossils of this group. Our analysis here, however, shows that this interpretation might not be the most parsimonious.

Specifically, we identify no signal of a super-archaic origin in any of the AMBN gene copies of Denisovans. The genomewide window-sliding analysis of all high coverage archaic genomes shows that AMBN falls within a region that bears no statistically significant signal for super-archaic ancestry. This is most evident when compared with a region of equal length that does, such as that around KRTAP-10. This result is also in agreement with previous genomewide scans of super-archaic ancestry in Denisovans and modern humans^35,36^, which did not identify AMBN as potentially super-archaic. Furthermore the phylogenetic analysis of the Denisova 3 haplotypes of AMBN, both bearing the rs564905233 SNP, places them closest to the Neanderthal haplotypes of the same gene. This placement is expected for the average Denisovan haplotype and is contrary to the expectations of a super-archaic one.

Our investigation of the Denisova 25 genome does confirm that the rs564905233 SNP is in a heterozygous state, along with multiple other SNPs on the gene of AMBN. While the increased heterozygosity might be evidence of introgression, our results indicate that, if anything, the origin of this possible introgression is likely to be from a Neanderthal or even modern human ancestor, rather than a super-archaic population. We excluded contamination or incomplete lineage sorting as the cause of this modern human affinity, due to the 100kb length of the observation and the overall low levels of contamination for Denisova 25, which is estimated to be around 3%^24^. We speculate that this region of the Denisova 25 genome has received substantial Neanderthal introgression, which is supported by the equal genetic distance of Neanderthals and Denisova 3 to Denisova 25, both upstream and downstream of AMBN (Extended Data Fig. 3). This Neanderthal ancestry could perhaps be bearing a smaller segment of modern-human-like DNA, from earlier out-of-Africa expansion(s)^37–39^. This hypothesis however, would need to be investigated on a genomewide level and is beyond the scope of the current work. For the purpose of this work, our investigation supports that in Denisova 25, both AMBN pseudohaplotypes also do not show signals of super-archaic origin.

For clarity, we would like to highlight that we do not contest the fact that Denisovans admixed with a super-archaic lineage, which could turn out to be the same population of *Homo erectus* as the one sampled palaeoproteomically by Fu et al. 2026. Our position is that the Denisovan AMBN gene, including the rs564905233 SNP and its protein product, is unlikely to be originating from a super-archaic source. This, however, means that either Denisovans did not inherit this gene from the sampled *Homo erectus* population, but somehow share it with them, or they did inherit it from the hominin population to which the six fossils belong, however this *Homo erectus* population is not as “super-archaic” as one might expect. Inspired by the figure from Fu et al. 2026 we present here a couple of alternative scenarios that we believe could explain the observed data (Fig. 1B-1E).

### Alternative Models

In the first two models (Fig. 1B, 1C), the 6 *Homo erectus* samples investigated by Fu et al. 2026 belong to a population that is the product of admixture between *Homo erectus* and Denisovans. As a result, the sampled population inherited and incorporated both the Denisovan and *Homo erectus* AMBN mutations, while maintaining the *Homo erectus* morphology. In these models, the 6 samples could either be *Homo erectus* with Denisovan admixture (Fig. 1B) or Denisovans with *Homo erectus* admixture (Fig. 1C), depending on the percentage of each ancestry. Given, however, that all 6 samples show no evidence of heterozygosity in the AMBN variants, this admixture is unlikely to be very recent to the sampling time (∼400kya). This would be further supported by the fact that the two amino acid variants are found on the same protein. If each one of them originated from a different population, their combined haplotype would require some time for recombination to occur.

The two mutations are 2,322 basepairs apart, and using the hapmap recombination map^40^, this region has a recombination rate of 5.42e-08 recombination events per basepair, per generation. This is nearly 5 times that of the genome-wide average, which is 1.45 recombination events per basepair per generation (Extended Data Fig. 3). Thus, the probability that there will be a recombination between these two SNPs after 250,000 years is already ∼75%, and 99% after a million years, assuming the regional recombination rate. If one assumes a genomewide recombination rate, the probabilities are lowered to 25% and 68% accordingly. In either case, one cannot exclude that these two mutations have recombined from two originally separate haplotypes and/or populations.

A very different model to the above two, is that of Fig. 1D. In this model, the 6 *Homo erectus* samples are not “*Homo erectus*” per se, but a different Denisovan population to that of the Altai mountains, which possess its own unique mutations, in addition to the previously identified Denisovan mutations. Previous genomic studies have supported the existence of multiple Denisovan lineages, almost as diverged as Denisovans and Neanderthals ^20^.

Members of this network of Denisova populations could have either, both or neither of the two mutations of AMBN, depending on the frequency of those mutations.

In support of this model, the enamel protein data recovered by Fu et al. did not show any archaic variants compared to present day humans, Neanderthals and Denisovans.

Previously published enamel samples, including *Homo antecessor*^41^ , *Homo naledi*^42^ and *Paranthropus robustus*^43^ all show at least one variant that is shared with great apes to the exclusion of present day humans, Neanderthals and Denisovans, consistent with a longer divergence time from our lineage. While the absence of such markers in *Homo erectus* enamel could be the result of the low number of informative sites that are recoverable through palaeoproteomics, it might alternatively be a hint of a more recent divergence of the sampled specimens to our lineage than any of these other hominins (*Homo antecessor, Homo naledi, Paranthropus robustus*). On the contrary to this model, morphological analyses have consistently placed some of these samples as being more divergent and belonging to a separate clade to those belonging to the ‘Denisovan’ clade such as Penghu, Harbin, Dali and others^2,44^.

Our last proposed model (Fig. 1E) is one where the observed affinity between the sampled *Homo erectus* specimens and the Denisovans is not due to an admixture event or a close genetic relationship. In this model either one or both mutations of AMBN occurred in the last common ancestor of the two populations. Later, due to incomplete lineage sorting, both Denisovans and these *Homo erectus* samples inherited the mutation in high frequencies, while modern humans and Neanderthals did not. The likelihood of this scenario heavily depends on the effective population size of the common ancestral population and its genetic diversity. A very large and genetically diverse ancestral population makes this a possible consideration, while a smaller population makes this random occurrence very unlikely.

### Resolving the puzzle

High quality genetic data could help resolve the above puzzle and point to one or more scenarios. The AMBN(A253G) mutation, found in these 6 *Homo erectus*, has so far not been recovered in any ancient DNA sample, or in any present day human within or outside an introgressed haplotype. Direct ancient DNA evidence, from a fossil bearing this mutation, would allow us to determine its demographic history. More recent fossils could possibly harbour this mutation, but recovery of DNA from the samples studied by Fu et al. 2026 is unlikely, given the age of the samples (∼400 ky). Alternatively, the recovery of this mutation in high quality genetic data from modern humans, would allow us to investigate whether the segment bearing the mutation shows a super-archaic profile or not. A positive signal would support models B and C, while a simple “Denisovan” signal would support models D or E. It is however still unclear if the AMBN(A253G) mutation will be found in any present day human.

Palaeoproteomics could also contribute to the unraveling of this puzzle. Future enamel sampling and the identification of the AMBN(A253G) variant, and not AMBN(A273V), in older samples, preferably with a clear *Homo erectus* morphology, would also support the B and C models. Given the recovery of hominin enamel proteins from samples as old as 2 to 3 million years^43,45^, tracking these mutations even in the oldest available fossils should be possible.

On the contrary, finding the AMBN(A253G) mutation alone, in late, morphologically “Denisovan” samples, would increase support for model D. However one would still be able to make the case of the variant presence in late Denisovans as the product of introgression. In any case, the identification of AMBN(A253G) can be used to target samples for ancient DNA sequencing. A different breakthrough could come from the recovery of dentine or bone proteins from this *Homo erectus* population. Given the higher number of proteins present in those tissues, one could observe whether this pattern of Denisovan affinity persists, and whether it coincides with additional “*Homo erectus*” mutations (mutations that have not been observed or that are closer to great apes than to modern humans). Unfortunately, faunal material tested by Fu et al. for dentin and bone preservation, did not yield positive results^22^.

Lastly, but equally important to the molecular analyses, morphological data could also help resolve this puzzle. A more in depth morphological analysis of the 6 teeth that have already been sampled, could help us understand whether their morphology could be described as clearly *Homo erectus*, with connections to earlier samples, especially African ones, or connections to the ever increasing amount of Denisovan material. Thanks to the enamel etching methodology employed by Fu et al, the 6 teeth should have their morphology preserved for further studies.

In conclusion, the Middle and Late Pleistocene hominin fossil record remains open to multiple interpretations in terms of population connections and taxonomy. Here, we use high quality genetic data to show that the amino acid variant found in Middle and Late Pleistocene Denisovan genomes, and shared with *Homo erectus* samples, is unlikely to be introgressed from a super-archaic source. We provide further context for AMBN, a gene which is turning into a very useful marker bridging fossils and their morphology, ancient and modern DNA and palaeoproteomic data. While we caution against the interpretation of the protein data presented by Fu et al. 2026, we greatly acknowledge the quality and importance of this type of data, and commend the original authors for delivering it to the community. We fully expect that additional protein data, like the one presented by Fu et al. 2026, will be instrumental in disentangling the taxonomic confusion of Middle and Late Pleistocene Asia when combined with morphological and DNA data.

## Supporting information

Supplementary Material Folder (Figures, Tables and Scripts)

## Data and Code Availability

All data and tools used for the analysis and visualisation are publicly available.

Code for assembling the FASTA dataset, calculating the genetic distances between humans, Neanderthals and Denisovans, plotting this distance and plotting the phylogenetic trees is written in BASH and Python 3 and is available in the Supplementary folder as 3 separate scripts.

---

**Extended Data Fig. 1.**
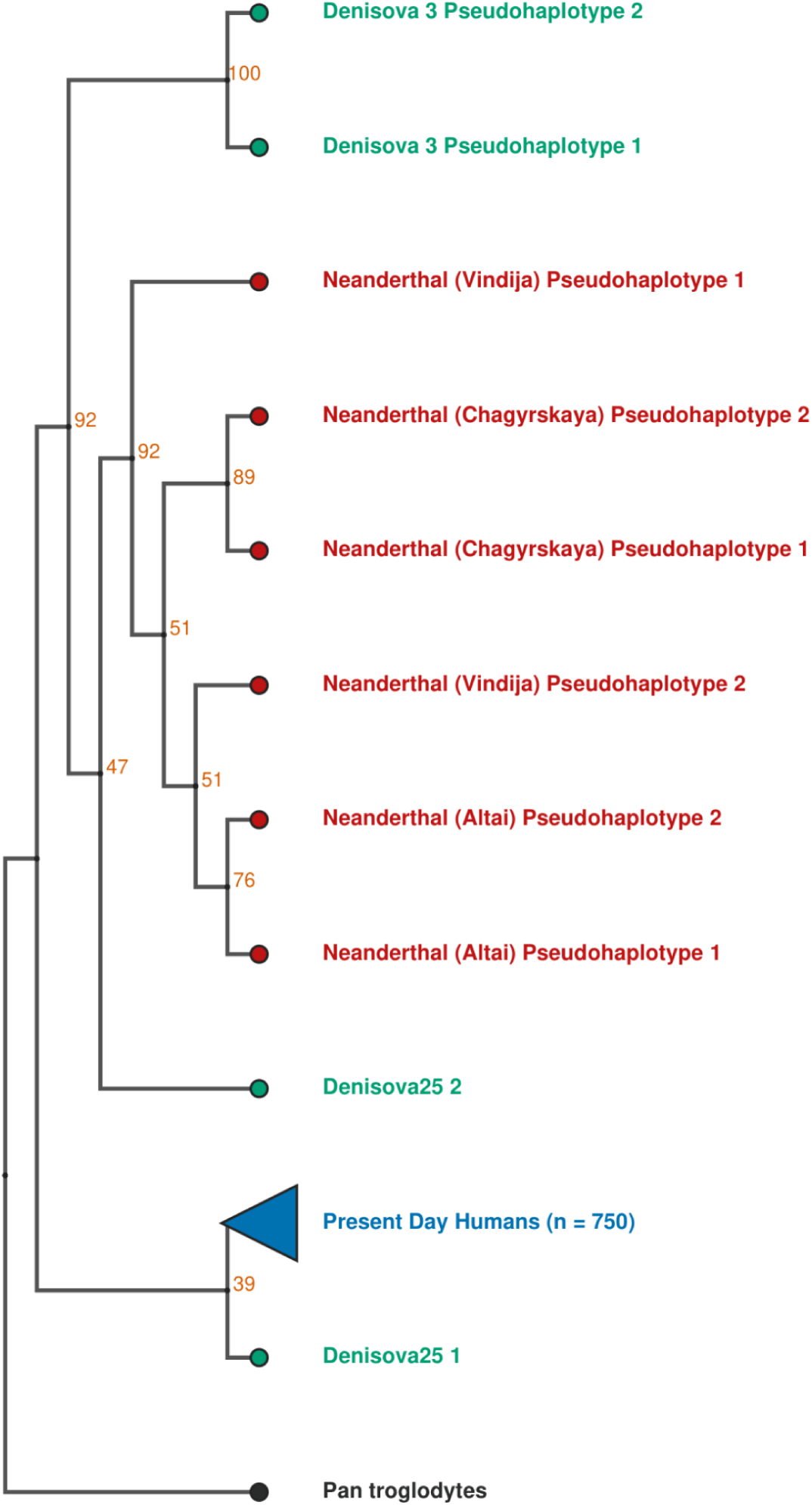
Phylogenetic tree of Denisova 25 AMBN hapotypes. Maximum likelihood phylogenetic tree of the AMBN haplotypes for Neanderthals, present day humans, Denisova 3 and Denisova 25. The tree is rooted using *Pan paniscus*. Bootstrap support values of each internal node are shown in orange.

**Extended Data Fig. 2.**
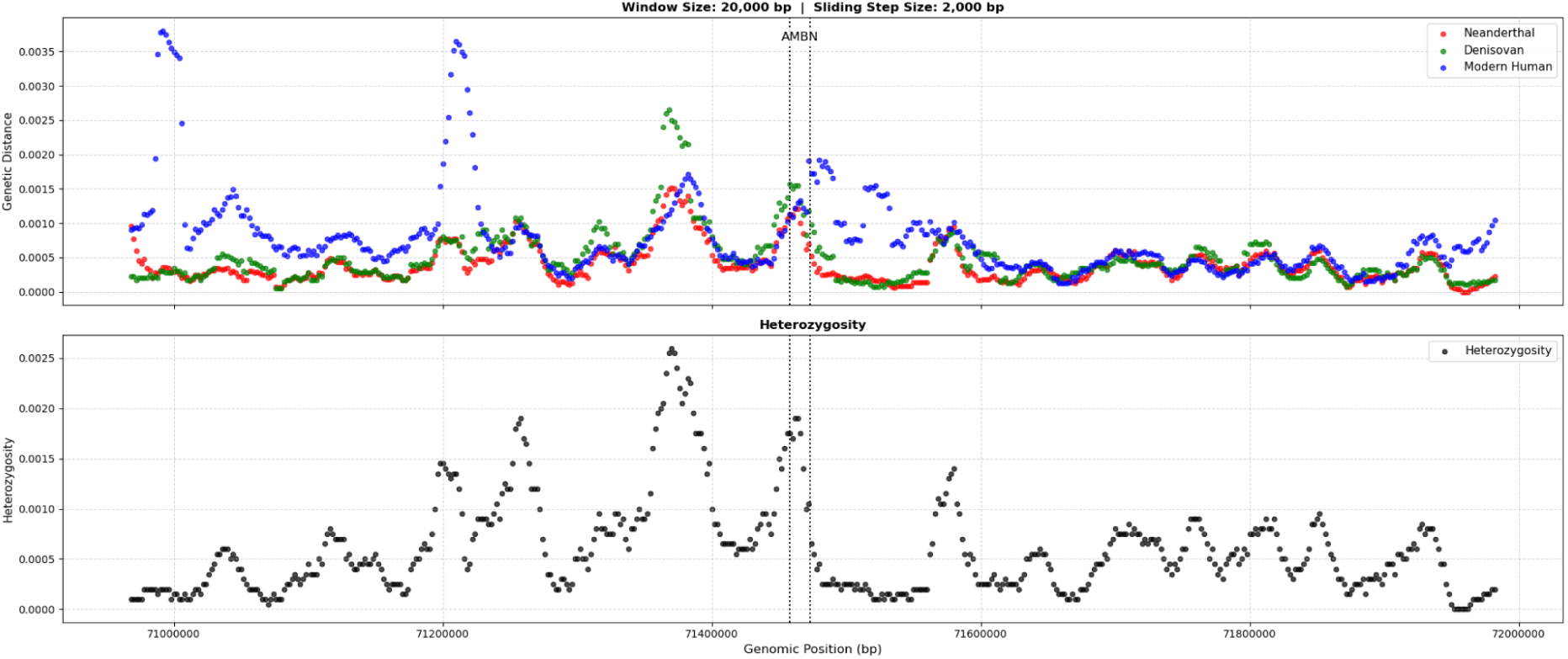
Sliding window analysis. Sliding window analysis showing the genetic distance of a group to Denisova 25 (top) and the heterozygosity of Denisova 25 (bottom). Each dot in the top plot represents the distance, calculated within a 20kb window (with a 2kb sliding step), from present day Africans (blue), Neanderthals (red) and Denisova 3 (green). Each dot in the bottom plot represents the heterozygosity of Denisova 25 in a 20kb window.

**Extended Data Fig. 3.**
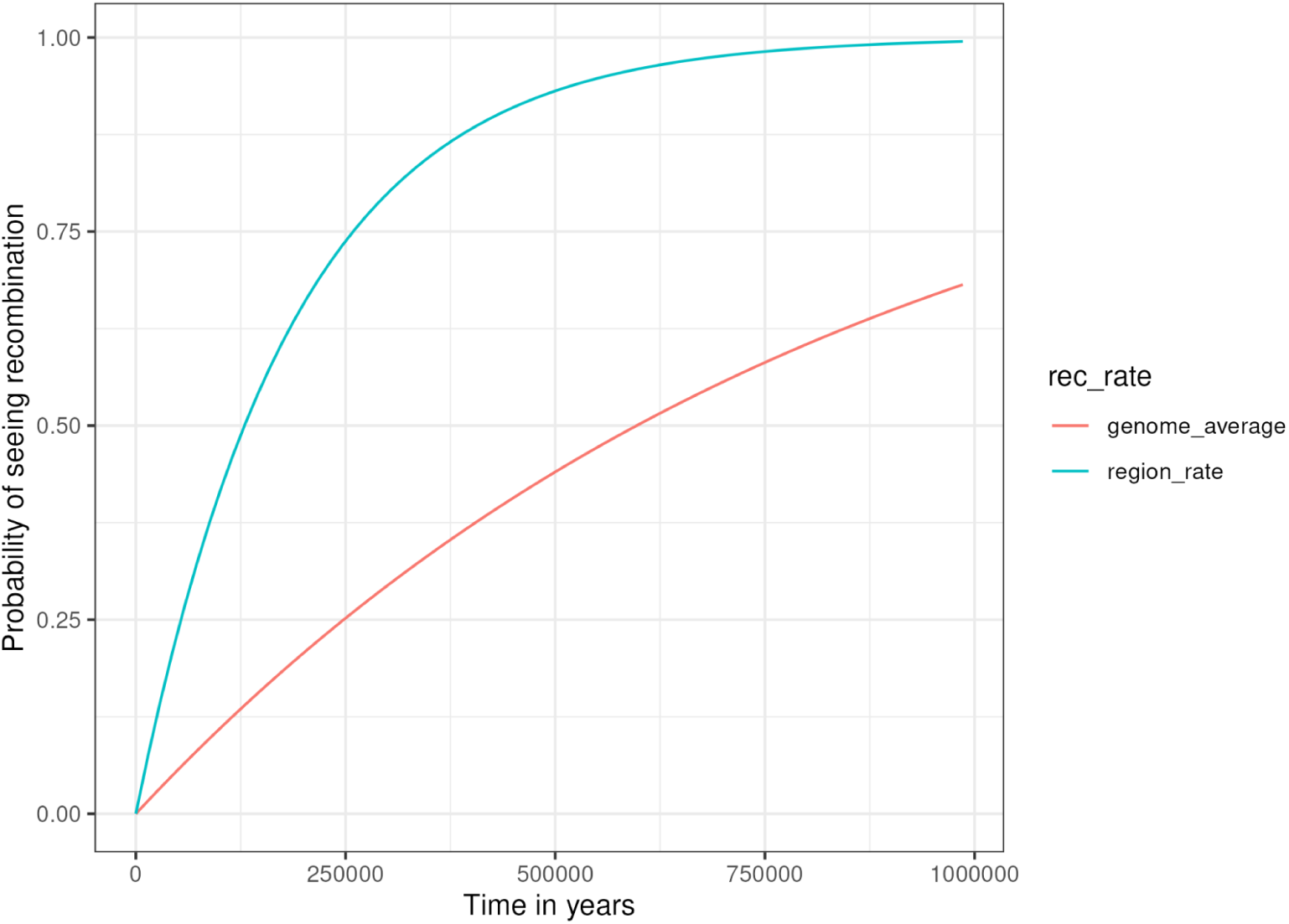
Recombination probability. Recombination probability over 1 million years given the recombination rate of the region (cyan) or the genomewide average (red).

**Extended Data Table 1.** AMBN Allelic states. Table recording the allelic state of the AMBN gene for Denisova 25, compared to 4 high coverage archaic individuals and the allele frequency of present day Africans from Gnomad. Allelic states of Denisova 25 highlighted with a dark outline were missing from the published VCF file and called by looking at the aligned reads of the bam file. Allelic frequencies of Africans highlighted by a dotted outline were missing from Gnomad callset completely and thus inferred to be close to 0.

| Position on the 4th Chromosome (Hg19) | Reference Allele | Alternative Allele | Altai Neanderthal (D5) | Vindija Neanderthal (33) | Chagyrskaya Neanderthal | Denisova 3 (D3) | Denisova 25 (D25) | Gnomad V4 Africa Frequency |
| --- | --- | --- | --- | --- | --- | --- | --- | --- |
| 71458018 | A | G | 1/1 | 1/1 | 1/1 | 1/1 | 0/1 | 0.0 |
| 71458331 | C | T | 1/1 | 1/1 | 1/1 | 1/1 | 0/1 | 0.0 |
| 71458399 | C | T | 1/1 | 1/1 | 1/1 | 1/1 | 0/1 | 0.0 |
| 71458546 | G | A | 1/1 | 1/1 | 1/1 | 1/1 | 0/1 | 0.00380356 |
| 71458732 | C | T | 1/1 | 1/1 | 1/1 | 1/1 | 0/1 | 0.786035 |
| 71458764 | A | T | 0/0 | 0/0 | 0/0 | 0/1 | 0/0 | 0.0 |
| 71458765 | C | T | 0/0 | 0/0 | 0/0 | 0/1 | 0/0 | 0.0 |
| 71458766 | A | T | 0/1 | 0/0 | 0/1 | 0/0 | 0/0 | 0.0 |
| 71458767 | A | T | 0/1 | 0/0 | 0/1 | 0/0 | 0/0 | 0.0 |
| 71458877 | T | C | 0/0 | 0/0 | 0/0 | 1/1 | 0/0 | 0.0 |
| 71458885 | G | A,- | 1/1 | 1/1 | 1/1 | 1/1 | 0/1 | 0.006921 |
| 71458939 | G | T | 1/1 | 1/1 | 1/1 | 1/1 | 0/1 | 0.0 |
| 71459289 | C | T | 1/1 | 1/1 | 1/1 | 1/1 | 0/1 | 0.0 |
| 71462894 | A | G | 1/1 | 1/1 | 1/1 | 0/0 | 0/1 | 0.0 |
| 71463009 | T | C | 1/1 | 1/1 | 1/1 | 1/1 | 0/1 | 0.0 |
| 71463052 | C | A | 1/1 | 1/1 | 1/1 | 1/1 | 0/1 | 0.0 |
| 71463074 | A | G,T | 1/1 | 1/1 | 1/1 | 1/1 | 0/1 | 0.0 |
| 71463436 | C | T | 1/1 | 1/1 | 1/1 | 1/1 | 0/1 | 0.000264945 |
| 71463489 | G | C | 1/1 | 1/1 | 1/1 | 1/1 | 0/1 | 0.0 |
| 71463537 | T | G | 1/1 | 1/1 | 1/1 | 1/1 | 0/1 | 0.0 |
| 71464007 | T | G,C | 1/1 | 1/1 | 1/1 | 1/1 | 0/1 | 0.0004266 |
| 71464229 | G | A | 0/0 | 0/0 | 0/0 | 1/1 | 0/0 | 0.0 |
| 71464276 | T | C | 1/1 | 1/1 | 1/1 | 1/1 | 0/1 | 0.0 |
| 71464290 | A | G | 1/1 | 0/1 | 0/0 | 0/0 | 0/0 | 0.0 |
| 71464312 | G | T | 1/1 | 1/1 | 1/1 | 1/1 | 0/1 | 0.0 |
| 71464464 | C | T | 1/1 | 1/1 | 1/1 | 1/1 | 1/1 | 0.791233 |
| 71465264 | C | T | 1/1 | 1/1 | 1/1 | 1/1 | 0/1 | 0.0 |
| 71465301 | G | A | 0/0 | 0/1 | 0/0 | 0/0 | 0/0 | 0.000169409 |
| 71465533 | T | C | 0/0 | 0/1 | 1/1 | 0/0 | 0/0 | 0.0 |
| 71466564 | T | G | 0/0 | 0/1 | 0/0 | 0/0 | 0/0 | 0.0 |
| 71466582 | A | C | 0/0 | 0/0 | 0/0 | 1/1 | 0/0 | 0.0 |
| 71466584 | C | A | 0/0 | 0/0 | 0/0 | 1/1 | 0/0 | 0.000192539 |
| 71466802 | T | C | 0/0 | 0/0 | 0/0 | 1/1 | 0/0 | 0.0 |
| 71466914 | C | T | 1/1 | 1/1 | 1/1 | 1/1 | ./. | 0.653408 |
| 71467443 | A | G | 1/1 | 1/1 | 1/1 | 1/1 | 0/1 | 0.0626563 |
| 71467593 | A | C | 1/1 | 1/1 | 1/1 | 1/1 | 1/1 | 0.0 |
| 71467732 | C | A | 1/1 | 1/1 | 1/1 | 0/0 | 0/1 | 0.186389 |
| 71468590 | A | T,G | 0/0 | 0/0 | 1/1 | 0/0 | 0/0 | 0.0 |
| 71469953 | C | T | 1/1 | 1/1 | 1/1 | 1/1 | 1/1 | 0.973601 |
| 71470174 | A | G | 1/1 | 1/1 | 1/1 | 1/1 | 0/1 | 0.166651 |
| 71470298 | G | A,T | 1/1 | 1/1 | 1/1 | 1/1 | 0/1 | 0.04141 |
| 71470419 | T | C | 1/1 | 1/1 | 1/1 | 1/1 | 0/1 | 0.216712 |
| 71471258 | G | T | 1/1 | 1/1 | 1/1 | 1/1 | 1/1 | 0.233182 |
| **71471920** | **A** | **G** | **0/0** | **0/0** | **0/0** | **1/1** | **0/1** | **0.0** |
| 71472426 | A | G | 1/1 | 1/1 | 1/1 | 1/1 | 1/1 | 0.590286 |
| 71472560 | C | T | 0/0 | 0/0 | 0/0 | 1/1 | ./. | 0.0925953 |

**\* All figures and tables are available in higher quality as separate files in the supplementary folder.**

## Acknowledgements

We would like to thank Viridiana Villa-Islas for helpful discussions during the analysis of the data.

