## Supplementary Material Folder (Figures, Tables and Scripts) for "No signal of a super-archaic origin of the Denisovan AMBN gene, a comment on the protein affinity of *Homo erectus* and Denisovan enamel proteins": Den_25_Summary_of_Findings.pdf

### AMBN of Denisova 25 and it's surounding region

A

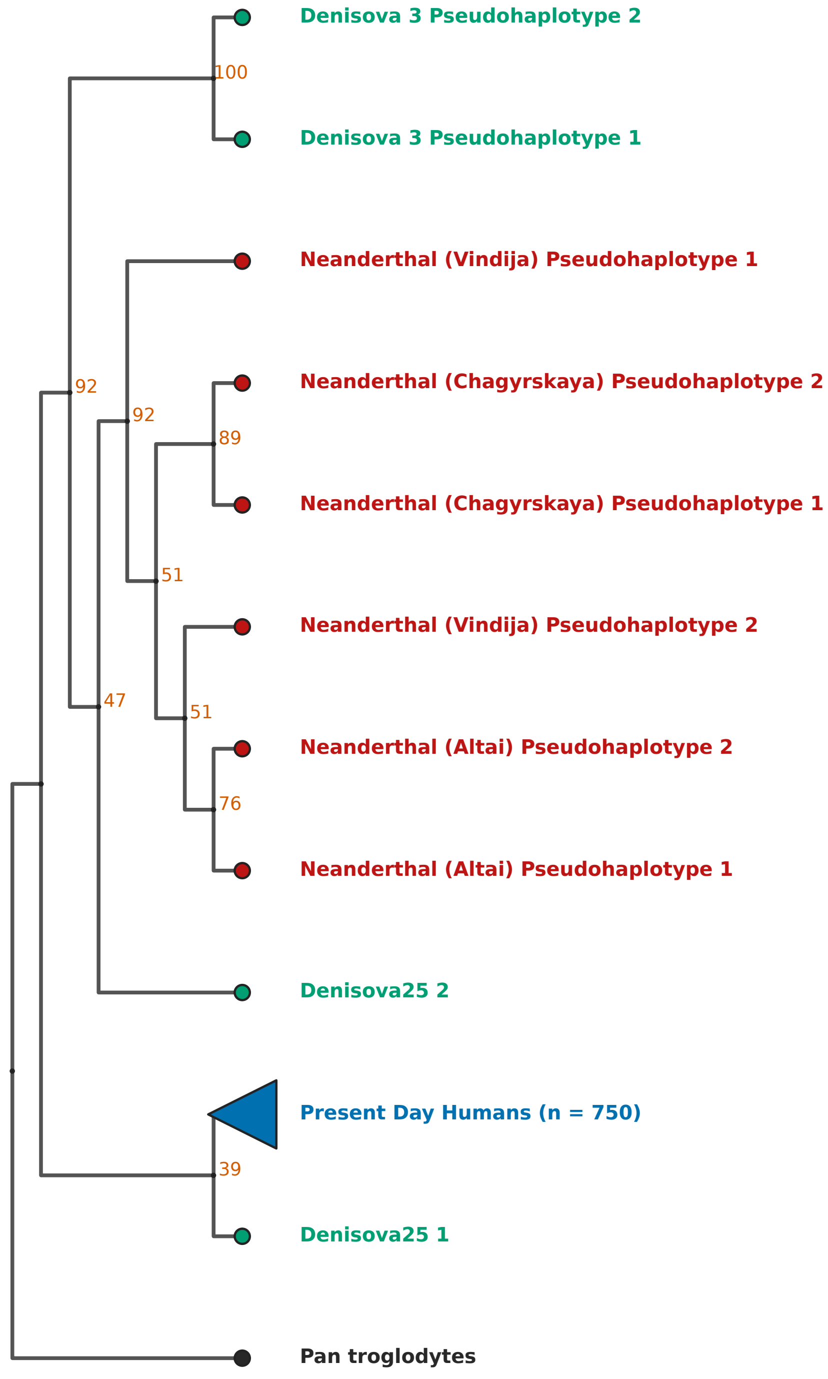


B

| Chromosome | Position | REF | ALT | Altai Neandertal ( | Vindija Neanderthal (33) | Chagyrskaya Neanderthal | Denisova 3 (D3 | Denisova 25 (D2 | Gnomad V4 Africa Frequency |
|---|---|---|---|---|---|---|---|---|---|
| 4 | 71458018 | A | G | 1/1 | 1/1 | 1/1 | 1/1 | 0/1 | 0.0 |
| 4 | 71458331 | C | T | 1/1 | 1/1 | 1/1 | 1/1 | 0/1 | 0.0 |
| 4 | 71458399 | C | T | 1/1 | 1/1 | 1/1 | 1/1 | 0/1 | 0.0 |
| 4 | 71458546 | G | A | 1/1 | 1/1 | 1/1 | 1/1 | 0/1 | 0.00380356 |
| 4 | 71458732 | C | T | 1/1 | 1/1 | 1/1 | 1/1 | 0/1 | 0.786035 |
| 4 | 71458764 | A | T | 0/0 | 0/0 | 0/0 | 0/1 | 0/0 | 0.0 |
| 4 | 71458765 | C | T | 0/0 | 0/0 | 0/0 | 0/1 | 0/0 | 0.0 |
| 4 | 71458766 | A | T | 0/1 | 0/0 | 0/1 | 0/0 | 0/0 | 0.0 |
| 4 | 71458767 | A | T | 0/1 | 0/0 | 0/1 | 0/0 | 0/0 | 0.0 |
| 4 | 71458877 | T | C | 0/0 | 0/0 | 0/0 | 1/1 | 0/0 | 0.0 |
| 4 | 71458885 | G | A,- | 1/1 | 1/1 | 1/1 | 1/1 | 0/1 | 0.006921 |
| 4 | 71458939 | G | T | 1/1 | 1/1 | 1/1 | 1/1 | 0/1 | 0.0 |
| 4 | 71459289 | C | T | 1/1 | 1/1 | 1/1 | 1/1 | 0/1 | 0.0 |
| 4 | 71462894 | A | G | 1/1 | 1/1 | 1/1 | 0/0 | 0/1 | 0.0 |
| 4 | 71463009 | T | C | 1/1 | 1/1 | 1/1 | 1/1 | 0/1 | 0.0 |
| 4 | 71463052 | C | A | 1/1 | 1/1 | 1/1 | 1/1 | 0/1 | 0.0 |
| 4 | 71463074 | A | G,T | 1/1 | 1/1 | 1/1 | 1/1 | 0/1 | 0.0 |
| 4 | 71463436 | C | T | 1/1 | 1/1 | 1/1 | 1/1 | 0/1 | 0.000264945 |
| 4 | 71463489 | G | C | 1/1 | 1/1 | 1/1 | 1/1 | 0/1 | 0.0 |
| 4 | 71463537 | T | G | 1/1 | 1/1 | 1/1 | 1/1 | 0/1 | 0.0 |
| 4 | 71464007 | T | G,C | 1/1 | 1/1 | 1/1 | 1/1 | 0/1 | 0.0004266 |
| 4 | 71464229 | G | A | 0/0 | 0/0 | 0/0 | 1/1 | 0/0 | 0.0 |
| 4 | 71464276 | T | C | 1/1 | 1/1 | 1/1 | 1/1 | 0/1 | 0.0 |
| 4 | 71464290 | A | G | 1/1 | 0/1 | 0/0 | 0/0 | 0/0 | 0.0 |
| 4 | 71464312 | G | T | 1/1 | 1/1 | 1/1 | 1/1 | 0/1 | 0.0 |
| 4 | 71464464 | C | T | 1/1 | 1/1 | 1/1 | 1/1 | 1/1 | 0.791233 |
| 4 | 71465264 | C | T | 1/1 | 1/1 | 1/1 | 1/1 | 0/1 | 0.0 |
| 4 | 71465301 | G | A | 0/0 | 0/1 | 0/0 | 0/0 | 0/0 | 0.000169409 |
| 4 | 71465533 | T | C | 0/0 | 0/1 | 1/1 | 0/0 | 0/0 | 0.0 |
| 4 | 71466564 | T | G | 0/0 | 0/1 | 0/0 | 0/0 | 0/0 | 0.0 |
| 4 | 71466582 | A | C | 0/0 | 0/0 | 0/0 | 1/1 | 0/0 | 0.0 |
| 4 | 71466584 | C | A | 0/0 | 0/0 | 0/0 | 1/1 | 0/0 | 0.000192539 |
| 4 | 71466802 | T | C | 0/0 | 0/0 | 0/0 | 1/1 | 0/0 | 0.0 |
| 4 | 71466914 | C | T | 1/1 | 1/1 | 1/1 | 1/1 | ./. | 0.653408 |
| 4 | 71467443 | A | G | 1/1 | 1/1 | 1/1 | 1/1 | 0/1 | 0.0626563 |
| 4 | 71467593 | A | C | 1/1 | 1/1 | 1/1 | 1/1 | 1/1 | 0.0 |
| 4 | 71467732 | C | A | 1/1 | 1/1 | 1/1 | 0/0 | 0/1 | 0.186389 |
| 4 | 71468590 | A | T,G | 0/0 | 0/0 | 1/1 | 0/0 | 0/0 | 0.0 |
| 4 | 71469953 | C | T | 1/1 | 1/1 | 1/1 | 1/1 | 1/1 | 0.973601 |
| 4 | 71470174 | A | G | 1/1 | 1/1 | 1/1 | 1/1 | 0/1 | 0.166651 |
| 4 | 71470298 | G | A,T | 1/1 | 1/1 | 1/1 | 1/1 | 0/1 | 0.04141 |
| 4 | 71470419 | T | C | 1/1 | 1/1 | 1/1 | 1/1 | 0/1 | 0.216712 |
| 4 | 71471258 | G | T | 1/1 | 1/1 | 1/1 | 1/1 | 1/1 | 0.233182 |
| **4** | **71471920** | **A** | **G** | **0/0** | **0/0** | **0/0** | **1/1** | **0/1** | 0.0 |
| 4 | 71472426 | A | G | 1/1 | 1/1 | 1/1 | 1/1 | 1/1 | 0.590286 |
| 4 | 71472560 | C | T | 0/0 | 0/0 | 0/0 | 1/1 | ./. | 0.0925953 |

Sites that have been called by looking at reads in the BAM file. ./. signidfies that there were too few reads to call

Absense from Gnomad, interpreted as 0.0 frequency

C

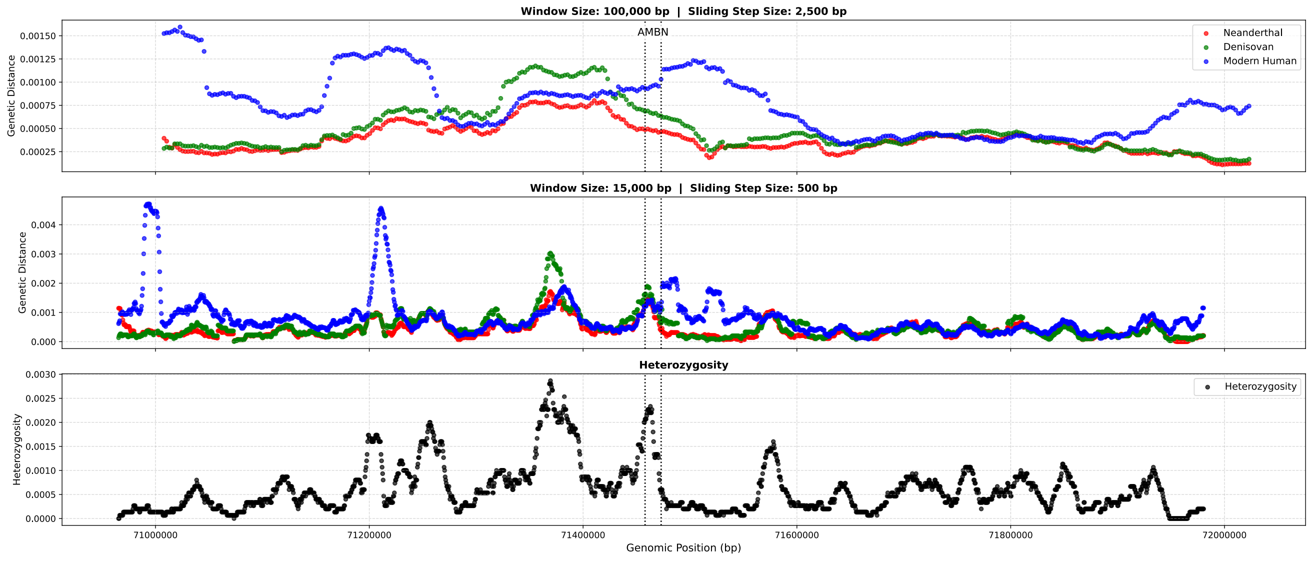
