## Supplementary Material Folder (Figures, Tables and Scripts) for "No signal of a super-archaic origin of the Denisovan AMBN gene, a comment on the protein affinity of *Homo erectus* and Denisovan enamel proteins": Count_Differences_Hg_19.py

```
import gzip
#### import vcfpy ### needs pip to install
import pysam ### from conda
import pysam.samtools
import pysam.bcftools
import math 
import statistics
from pysam import VariantFile


################################################################################################################################################################
############## Functions

##### Returns two dictionaries, one with alleles of the 5 high coverage archaics and one with the frequencies of the alleles for Africans. The two dicitonaries share the same sites.
def Get_States_of_Fixed_All_Samples(CHR, Start, End):

    ########## Loop analysis

    ###  High Coverage Archaics
    Archaic_Genomes = F'/projects/archaic_introgression-AUDIT/data/resources/Archaic_genomes/hg19_vcf/individuals_highcov.{CHR}.bcf'
    Archaic_Genomes = F'/projects/archaic_introgression-AUDIT/data/TEMP_for_Herectus/Archaics_High_Coverage/Archaics_BCF_Zenodo/highcov_ind_{CHR}.bcf'

    ### Den25 High Coverage
    Den_25_Genome = F"/projects/archaic_introgression-AUDIT/data/TEMP_for_Herectus/Archaics_High_Coverage/Den25/chr{CHR}.Den25.L35MQ25.B30.map35_100.vcf.gz"

    ### Gnomad modern humans
    Gnomad_Genomes = F"/projects/archaic_introgression-AUDIT/data/TEMP_for_Herectus/Modern_Humans/gnomad.genomes.r2.1.1.sites.{CHR}.vcf.bgz"


    ###################################

    ############# Step 1 - Load VCF/BCF files

    Archaics_BCF = VariantFile(Archaic_Genomes)
    Den25_VCF = VariantFile(Den_25_Genome)
    Moderns_VCF = VariantFile(Gnomad_Genomes)


    ##### First get positions where 4 high coverage archaics are fixed

    Region_variants_Archaic = list(Archaics_BCF.fetch(F"{CHR}",Start,End))

    Pos_of_interest = {}

    for VAR in Region_variants_Archaic:
        
        Den_3 = [x for x in VAR.samples["Denisova"].allele_indices ]
        Nea_Alt = [x for x in VAR.samples["AltaiNeandertal"].allele_indices ]
        Nea_Vind = [x for x in VAR.samples["Vindija33.19"].allele_indices ]
        Nea_Chag = [x for x in VAR.samples["Chagyrskaya-Phalanx"].allele_indices ]
        

        Pos_of_interest[int(VAR.pos)] = {"Denisova_3":Den_3, "Altai_Neandertal":Nea_Alt , "Vindija_Neanderthal":Nea_Vind , "Chagyrskaya_Neanderthal":Nea_Chag}


    ##### Then, see what modern humans from Africa have for same region

    Region_variants_Modern= list(Moderns_VCF.fetch(F"{CHR}",Start,End)) 
    Pos_Afr_Freq = {}

    for VAR in Region_variants_Modern:
        
        Position = int(VAR.pos)
        
        #### For positions where archaics have variants
        if (Position in Pos_of_interest.keys()) and (Position not in Pos_Afr_Freq.keys()): ### avoid adding the same position twice (triallelic)
            
            if "AF_afr" in VAR.info.keys():
                African_Freq = float(VAR.info['AF_afr'][0]) ### Its a tupple, with first item being the frequency
            else:
                African_Freq = 0
                
            
            Pos_Afr_Freq[VAR.pos] = African_Freq
        


    ##### What to do if Gnomad did not cover that position     
    for Position in Pos_of_interest.keys():        
        if Position not in Pos_Afr_Freq.keys():
            Pos_Afr_Freq[Position] = 0


    #### For positions of interest, add Denisova 25
        
    Region_variants_Den_25 = list(Den25_VCF.fetch(F"{CHR}",Start,End)) 


    for VAR in Region_variants_Den_25:
        
        Position = int(VAR.pos)
        
        if Position in Pos_of_interest.keys():
          
            Den_25 = [x for x in VAR.samples["Denisova25"].allele_indices ]
            Pos_of_interest[VAR.pos]["Denisova_25"] = Den_25


    return Pos_of_interest, Pos_Afr_Freq


########## Distance Calculator

def Calculate_Distance_from_Den25(Dic1, Dic2):
    
    
    Neanderthal_Distance = []
    Denisova_Distance = []
    Africans_Distance = []
    

    Counted_Pos = 0
    Heterozygous_Pos = 0

    for POS in Dic1.keys():
        
        for J,K in Dic1[POS].items():
            if K == [None, None]:
                Dic1[POS][J]= [0,0]
        
        if Dic2[POS] <= 0.1:
            Dic2[POS]=0
        
        if "Denisova_25" not in Dic1[POS].keys():
            continue
        
        Neanderthal_Freq = statistics.mean([ y for x in [ Dic1[POS]["Altai_Neandertal"] , Dic1[POS]["Vindija_Neanderthal"], Dic1[POS]["Chagyrskaya_Neanderthal"]] for y in x ])
        Denisova_Freq = statistics.mean(Dic1[POS]["Denisova_3"])
        Africans_Freq = Dic2[POS]
        Den25_Freq = statistics.mean(Dic1[POS]["Denisova_25"])
        
        Counted_Pos+=1
        
        Neanderthal_Distance.append(abs(Den25_Freq - Neanderthal_Freq))
        Denisova_Distance.append(abs(Den25_Freq - Denisova_Freq))
        Africans_Distance.append(abs(Den25_Freq - Africans_Freq))
        
        if Den25_Freq == 0.5:
            Heterozygous_Pos+=1
    
    
    
    if Counted_Pos!=0:
        return sum(Neanderthal_Distance), sum(Denisova_Distance), sum(Africans_Distance), Heterozygous_Pos
    else:
        return 0, 0, 0, 0


#############################################################################################################################################################################################################################################
#### End of Functions


######## Chromosome
CHR = 4


##### For AMBN, "4",71457973,71473005 how does the region around it look ike
Max_Start = 71457973 - 500000
Max_End = 71473005 + 500000
Range = 100000
Step = 2500


Output_File = open(F"Differences_{CHR}_{Max_Start}_{Max_End}_{Range}kb_{Step}kbStep.txt",'w')
Output_File.write("Position\tDist_Nea\tDist_Den\tDist_AMH\tHeterozygocity\n")

for Start in range(Max_Start, Max_End, Step):
    
    
    End = Start + Range


    ###### get states of all 3 groups for this region
    Dic1, Dic2 = Get_States_of_Fixed_All_Samples(CHR, Start, End)
    ###### Calculate distances between groups and the Den 25
    Neanderthal_Distance, Denisova_Distance, Africans_Distance, Heterozygous_Value = [ x/(End-Start) for x in Calculate_Distance_from_Den25(Dic1, Dic2)]
    # print(Neanderthal_Distance, Denisova_Distance, Africans_Distance, Heterozygous_Value)
    Output_File.write(F"{CHR}:{Start}-{End}\t{Neanderthal_Distance}\t{Denisova_Distance}\t{Africans_Distance}\t{Heterozygous_Value}\n")
```
