## Supplementary Material Folder (Figures, Tables and Scripts) for "No signal of a super-archaic origin of the Denisovan AMBN gene, a comment on the protein affinity of *Homo erectus* and Denisovan enamel proteins": Extract_Haplotypes.sh

```
#!/bin/bash

#### HG19
###AMBN in hg19 4:71457973-71473005 

#### GRCH28
###AMBN in chr38 4:70592256-70607288 

### AMBN #Den Mutation# M-->V in GrCh38 4:70606203 


### AMBN #Herectus Mutation# A-->G in GrCh38 4:70603880-4:70603883

##### needed tools ### Need to be installed or loaded (conda should work too)
module load samtools
module load gsl/2.5
module load perl
module load bcftools


###### Create Fasta Dataset

read -r NAME CHR START END < Location.txt;

#### This is specific to Mjolnir server
samtools faidx ../RefGenome/hg19_refgenome.fa $CHR:$START-$END -c > $NAME.fa;

#### This is specific to Mjolnir server, Place the location of the vcfs/bcfs you want to extract all haplotypes
FILES = ../Differences_Counting/Archaics_BCF_Zenodo/ ../Differences_Counting/Den25;

for FILE in FILES
do
        bcftools query -l $FILE > Samples_This_VCF.txt;
        while read SAMPLE; do

                cat ./AMBN_Full.fa | bcftools consensus --missing ? -a ? -H 1 -s "$SAMPLE" -o ./Full_Gene/AMBN_"$SAMPLE"_1.fa "$FILE";
                fasta_file="AMBN_${SAMPLE}_1.fa";
                name="${fasta_file%.fa}";
                sed -i "1s|^>.*|>${name}|" "./Full_Gene/AMBN_${SAMPLE}_1.fa";

                cat ./AMBN_Full.fa | bcftools consensus --missing ? -a ? -H 2 -s "$SAMPLE" -o ./Full_Gene/AMBN_"$SAMPLE"_2.fa "$FILE";
                fasta_file="AMBN_${SAMPLE}_2.fa";
                name="${fasta_file%.fa}";
                sed -i "1s|^>.*|>${name}|" "./Full_Gene/AMBN_${SAMPLE}_2.fa";


        done<Samples_This_VCF.txt;

done;


###### Combine Data into one fasta file, align and run phylogenetic tree
#### Chimp Fasta file already downloaded and in Full_Gene folder from: https://www.ensembl.org/Pan_troglodytes/Gene/Sequence?db=core;g=ENSPTRG00000023351;r=4:60022649-60037696;t=ENSPTRT00000045270
module load mafft
module load iqtree2

cat ./Full_Gene/*.fa > ./Full_Gene/Combined/Combined_AMBN_Pan.fa;
mafft --auto ./Full_Gene/Combined/Combined_AMBN_Pan.fa > ./Full_Gene/Combined/Combined_AMBN_Pan_Aligned.fa;

#### IQ tree
iqtree2 -s ./Full_Gene/Combined/Combined_AMBN_Pan_Aligned.fa -m GTR+I -b 10;
```
