## Supplementary Material Folder (Figures, Tables and Scripts) for "No signal of a super-archaic origin of the Denisovan AMBN gene, a comment on the protein affinity of *Homo erectus* and Denisovan enamel proteins": Plot_Differences.py

```
import sys
import os
import matplotlib.pyplot as plt
import pandas as pd


#### Load provided file
Files = sys.argv[1:]

##### Define the region of interest (here AMBN)
region_start = 71457973
region_end = 71473005
region_mid = (region_start + region_end) / 2

##### Total rows = 1 per distance file + 1 extra row at the bottom for Heterozygocity
nrows = len(Files) + 1
fig, axes = plt.subplots(nrows=nrows, ncols=1, figsize=(21, 9), sharex=True)

###### Store dataframe of the last file to reuse for the Heterozygocity plot
last_df = None

######  Plot Genetic Distances for each file in axes[:-1]
for ax, file_name in zip(axes[:-1], Files):
    
    #### Error cacthing
    if not os.path.exists(file_name) or os.path.getsize(file_name) == 0:
        ax.text(0.5, 0.5, f"File missing or empty: {file_name}", 
                ha='center', va='center', transform=ax.transAxes)
        continue
    #### panda dataframe load file
    df = pd.read_csv(file_name, sep=r'\s+', on_bad_lines='skip', skip_blank_lines=True)
    df.dropna(subset=['Dist_Nea', 'Dist_Den', 'Dist_AMH', 'Heterozygocity'], inplace=True)
        
    if df.empty:
        ax.text(0.5, 0.5, f"No valid data in: {file_name}", 
                ha='center', va='center', transform=ax.transAxes)
        continue

    ##### Extract coordinates as numeric values (1st column)
    coords = df['Position'].str.extract(r':(\d+)-(\d+)').astype(int)
    df['Pos_Midpoint'] = (coords[0] + coords[1]) / 2

    ##### Save a reference if this is the last file
    if file_name == Files[-1]:
        last_df = df.copy()

    ###### Window length & Step size
    window_size = coords[1].iloc[0] - coords[0].iloc[0]
    step_size = (coords[0].iloc[1] - coords[0].iloc[0]) if len(coords) > 1 else 0

    ##### Scatter plot for distances (handdles 3 populations, change if more)
    ax.scatter(df['Pos_Midpoint'], df['Dist_Nea'], color='red', label='Neanderthal', s=20, alpha=0.7)
    ax.scatter(df['Pos_Midpoint'], df['Dist_Den'], color='green', label='Denisovan', s=20, alpha=0.7)
    ax.scatter(df['Pos_Midpoint'], df['Dist_AMH'], color='blue', label='Modern Human', s=20, alpha=0.7)

    ##### Vertical dotted lines for AMBN region
    ax.axvline(x=region_start, color='black', linestyle=':')
    ax.axvline(x=region_end, color='black', linestyle=':')

    ##### Subplot Heading
    title_text = f"Window Size: {window_size:,} bp  |  Sliding Step Size: {step_size:,} bp"
    ax.set_title(title_text, fontsize=12, fontweight='bold')
    ax.set_ylabel('Genetic Distance', fontsize=11)
    ax.grid(True, linestyle='--', alpha=0.5)


##### Add the plot of Heterozygocity for the LAST file in the final bottom subplot (axes[-1])
het_ax = axes[-1]

if last_df is not None and not last_df.empty:
    het_ax.scatter(last_df['Pos_Midpoint'], last_df['Heterozygocity'], color='black', label='Heterozygosity', s=20, alpha=0.7)
    het_ax.axvline(x=region_start, color='black', linestyle=':')
    het_ax.axvline(x=region_end, color='black', linestyle=':')
    het_ax.set_title(f"Heterozygosity", fontsize=12, fontweight='bold')
    het_ax.set_ylabel('Heterozygosity', fontsize=11)
    het_ax.grid(True, linestyle='--', alpha=0.5)
    het_ax.legend(fontsize=11, loc='upper right')
else:
    het_ax.text(0.5, 0.5, f"No valid Heterozygosity data in: {Files[-1]}", 
                ha='center', va='center', transform=het_ax.transAxes)


###### Add the "AMBN" text to the top subplot
axes[0].text(x=region_mid, y=0.95, s='AMBN', 
             transform=axes[0].get_xaxis_transform(),
             fontsize=12, ha='center', va='top', 
             bbox=dict(facecolor='white', alpha=0.8, edgecolor='none', pad=3))

##### X-axis formatting on the very bottom subplot
axes[-1].set_xlabel('Genomic Position (bp)', fontsize=12)
axes[-1].ticklabel_format(style='plain', axis='x')

##### Distance legend on top subplot
axes[0].legend(fontsize=11, loc='upper right')


#### OUput
plt.tight_layout()
plt.savefig("Plotted_Distances_Den_25.pdf")
plt.savefig("Plotted_Distances_Den_25.svg")
plt.savefig("Plotted_Distances_Den_25.png")
```
