## Supplementary Material Folder (Figures, Tables and Scripts) for "No signal of a super-archaic origin of the Denisovan AMBN gene, a comment on the protein affinity of *Homo erectus* and Denisovan enamel proteins": Tree_Plotting.py

```
import sys
import toytree
import toyplot.pdf


##### To swap names as they are in the fasta and .tree files
custom_name_mapping = {
    "Pan_tro_3_AMBN": "<i>Pan troglodytes</i>",
    "AMBN_AltaiNeandertal_1": "Neanderthal (Altai) Pseudohaplotype 1",
    "AMBN_AltaiNeandertal_2": "Neanderthal (Altai) Pseudohaplotype 2",
    "AMBN_Vindija33.19_1": "Neanderthal (Vindija) Pseudohaplotype 1",
    "AMBN_Vindija33.19_2": "Neanderthal (Vindija) Pseudohaplotype 2",
    "AMBN_Chagyrskaya-Phalanx_1": "Neanderthal (Chagyrskaya) Pseudohaplotype 1",
    "AMBN_Chagyrskaya-Phalanx_2": "Neanderthal (Chagyrskaya) Pseudohaplotype 2",
    "AMBN_Denisova_1": "Denisova 3 Pseudohaplotype 1",
    "AMBN_Denisova_2": "Denisova 3 Pseudohaplotype 2",
    "AMBN_Den_25_1": "Denisova 25 Pseudohaplotype 1",
    "AMBN_Den_25_2": "Denisova 25 Pseudohaplotype 2"}

#### User provided Tree file
INPUT_TREE = sys.argv[1]

#####Increase 'recursion limit' for big trees, will crash if not
sys.setrecursionlimit(50000)

##### load the tree and root it to pan
tree = toytree.tree(INPUT_TREE)
rtree = tree.root("Pan_tro_3_AMBN")


#### collapse clades consisting ONLY of target prefixes (e.g. modern humans)
target_prefixes = ("AMBN_NA", "AMBN_HG", "AMBN_HGDP")

clades_to_collapse = []
collapsed_leaves = set()

##### traverse tree
for node in rtree.traverse("preorder"):
    if not node.is_leaf(): ### leafs
        leaves = node.get_leaves()
        if any(leaf.name in collapsed_leaves for leaf in leaves): ###
            continue
        if all(leaf.name.startswith(target_prefixes) for leaf in leaves): #### if all leaves here are humans check this node to be collapsed (list/set)
            clades_to_collapse.append(node)
            for leaf in leaves:
                collapsed_leaves.add(leaf.name)

####### Drop tips (collapse nodes) that are in node list
tips_to_drop = []
rename_dict = {}

#### Cycle through nodes (clades) to be collapsed
for clade in clades_to_collapse:
    leaves = clade.get_leaves()
    rep_tip = leaves[0].name
    rep_tip = rep_tip.split("/")[0]
    rename_dict[rep_tip] = f"Present Day Humans (n = {len(leaves)})" ### Replace name with "present day ..."
    for leaf in leaves[1:]:
        tips_to_drop.append(leaf.name) #### for next step

#### Drop tips
pruned_tree = rtree.mod.drop_tips(*tips_to_drop)


### Plotting stuff

###### nicer colours
COLOR_OUTGROUP   = "#2B2B2B"  # Soft Charcoal / Black
COLOR_NEANDERTAL = "#BF1515"  # Red
COLOR_DENISOVA   = "#009E73"  # Green
COLOR_HUMAN      = "#0072B2"  # Blue

tip_labels = pruned_tree.get_tip_labels()
formatted_tip_labels = []
colors = []
tip_markers = []
tip_marker_sizes = []


##### format and replace tip labels with nicer labels and colour them
for tip in tip_labels:
    #######
    
    tip = tip.split("/")[0]
    
    if tip in rename_dict:
        display_name = rename_dict[tip]
    elif tip in custom_name_mapping:
        display_name = custom_name_mapping[tip]        
        
    else:
        #### Replaces raw file underscores with clean spaces for presentation
        display_name = tip.replace("AMBN_", "").replace("_", " ")
        
    formatted_tip_labels.append(display_name)
    

    ########## Check if tip in renamed dict
    is_collapsed = tip in rename_dict
    if is_collapsed:
        tip_markers.append("<")
        tip_marker_sizes.append(45)  ##### Slightly tuned marker size
    else:
        tip_markers.append("o")
        tip_marker_sizes.append(10)

    ########## add colour for this tip
    if "Pan_tro" in tip:
        colors.append(COLOR_OUTGROUP)
    elif any(neandertal in tip for neandertal in ["AltaiNeandertal", "Vindija", "Chagyrskaya"]):
        colors.append(COLOR_NEANDERTAL)
    elif "Denisova" in tip: 
        colors.append(COLOR_DENISOVA)
    elif is_collapsed or any(modern in tip for modern in ["NA", "HG", "HGDP"]):
        colors.append(COLOR_HUMAN)
    else:
        colors.append(COLOR_OUTGROUP) ## backup


#### For node labels
for node in pruned_tree.traverse():
    if node.is_leaf():
        node.display_support = ""  ### Hide on tips
    else:
        ##### Check if support exists and is valid
        if node.support is not None:
            try:
                ###### Convert to integer and store as a string in our new feature
                node.display_support = str(int(float(node.support)))
            except ValueError:
                node.display_support = ""
        else:
            node.display_support = ""


##### Tree Drawing!
canvas, axes, mark = pruned_tree.draw(
    ts = "s",                          ##### 'c' = Cladogram / Coalescent style (clean lines)   's' , 'r'
    tip_labels = formatted_tip_labels,
    tip_labels_colors = colors,
    tip_labels_align = True,
    tip_labels_style = {
        "anchor-shift": "38px",     # Clean gap between triangle and text
        "font-size": "13px",        # Slightly enlarged text
        "font-family": "sans-serif",# Clean modern font
        "font-weight": "bold"
    },
    edge_widths = 2.5,                 # Thicker, bolder branch lines
    edge_colors = "#555555",           # Neutral dark gray for inner connections
    width = 850,
    height = 1000,
    node_sizes = 2,
    node_colors = "black",
    node_labels = "display_support",
    node_labels_style={
        "font-size": "12px",          # Keep bootstrap text small
        "font-family": "sans-serif",
        "fill": "#D55E00",         # A nice color (like Vermillion) to make them pop
        "text-anchor": "start",     # Anchors text to the left so it flows right
        "anchor-shift": "10px",      # Pushes the text 4 pixels to the right
        "baseline-shift": "3px"
    
    },
    scale_bar = False                   # Adds scale bar at the bottom
)

##### Scalebar control
axes.x.ticks.show = True
axes.x.ticks.labels.location = "below"  # Moves the numbers above the scale bar line
axes.x.ticks.location = "below"

axes.x.ticks.style["transform"] = "scale(-1, 1)"


###### markers control
pruned_tree.annotate.add_tip_markers(
    axes = axes,
    marker = tip_markers,
    size = tip_marker_sizes,
    color = colors
)

##### output
toyplot.pdf.render(canvas, F"{INPUT_TREE}_Plotted_Tree.pdf")
toyplot.svg.render(canvas, F"{INPUT_TREE}_Plotted_Tree.svg")
```
