## Supplementary figures and images for "No signal of a super-archaic origin of the Denisovan AMBN gene, a comment on the protein affinity of *Homo erectus* and Denisovan enamel proteins"

### Fig1_extnd.pdf

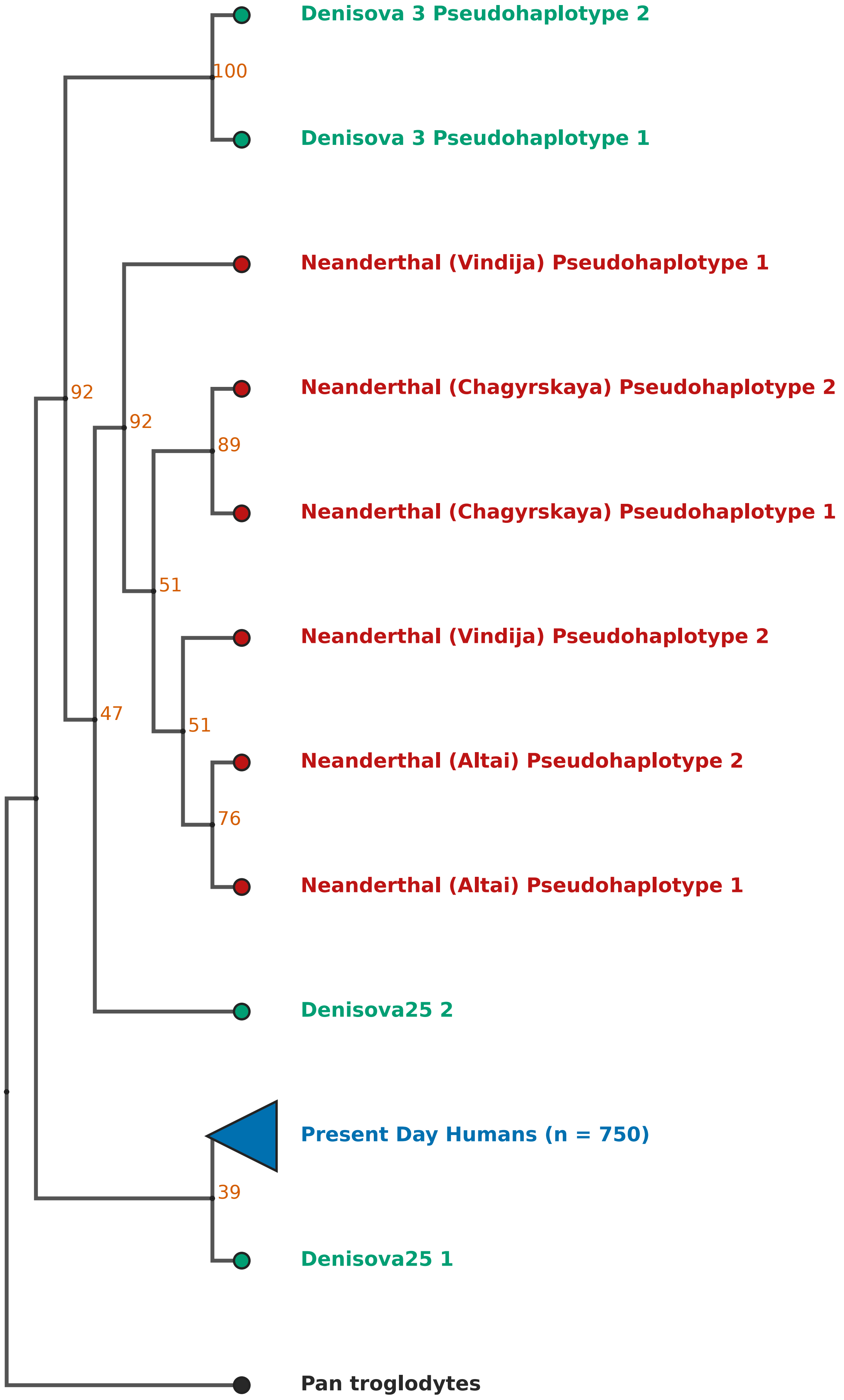

### Fig1_main.pdf

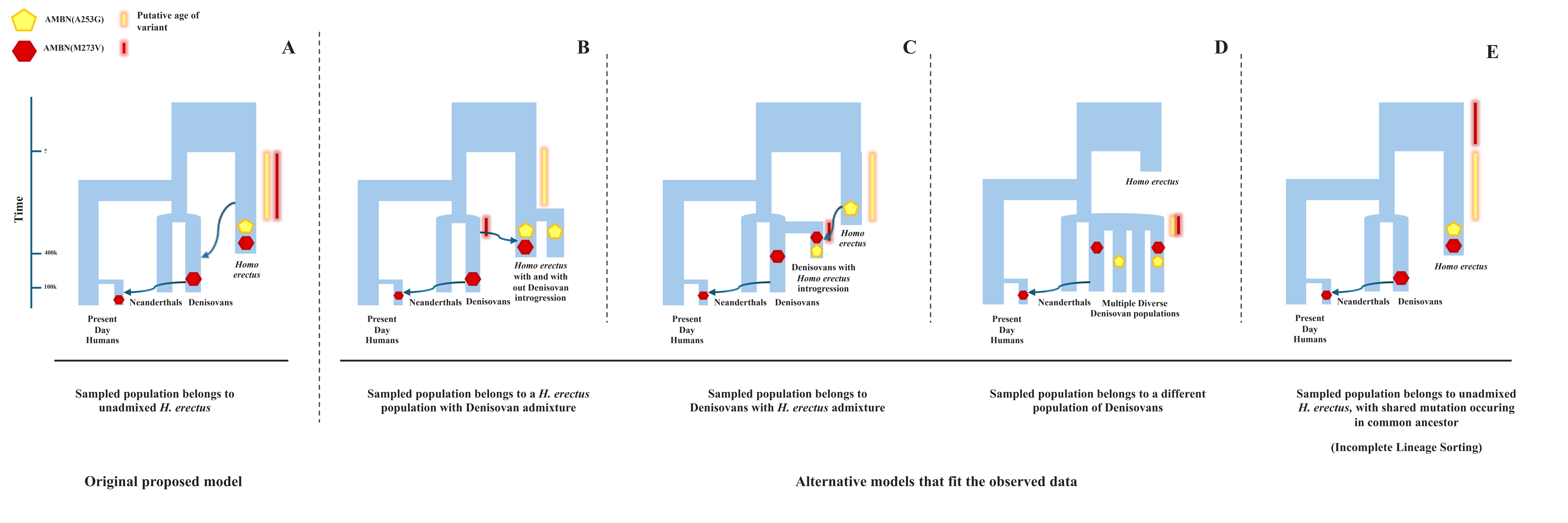

### Fig2_extnd.pdf

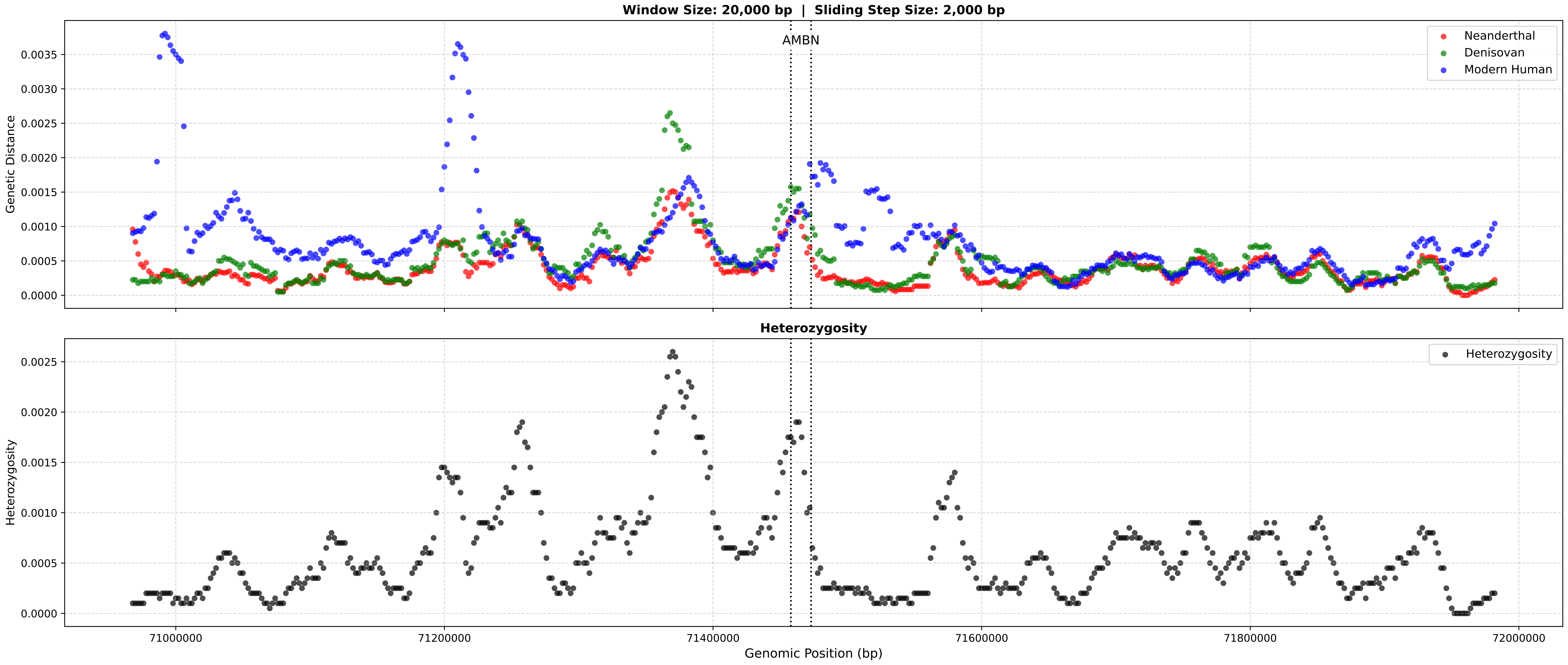

### Fig2_main.pdf

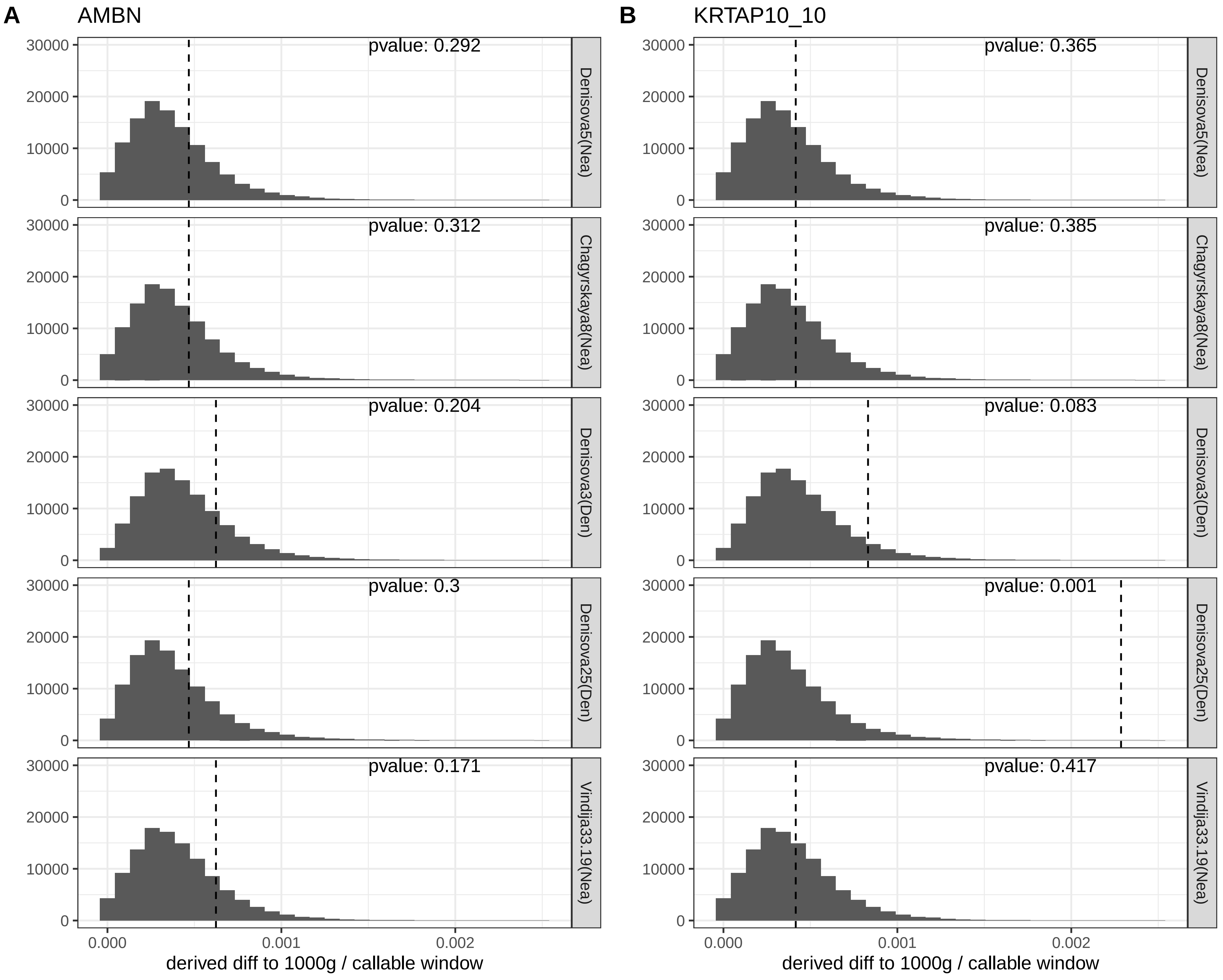

### Fig3_extnd.pdf

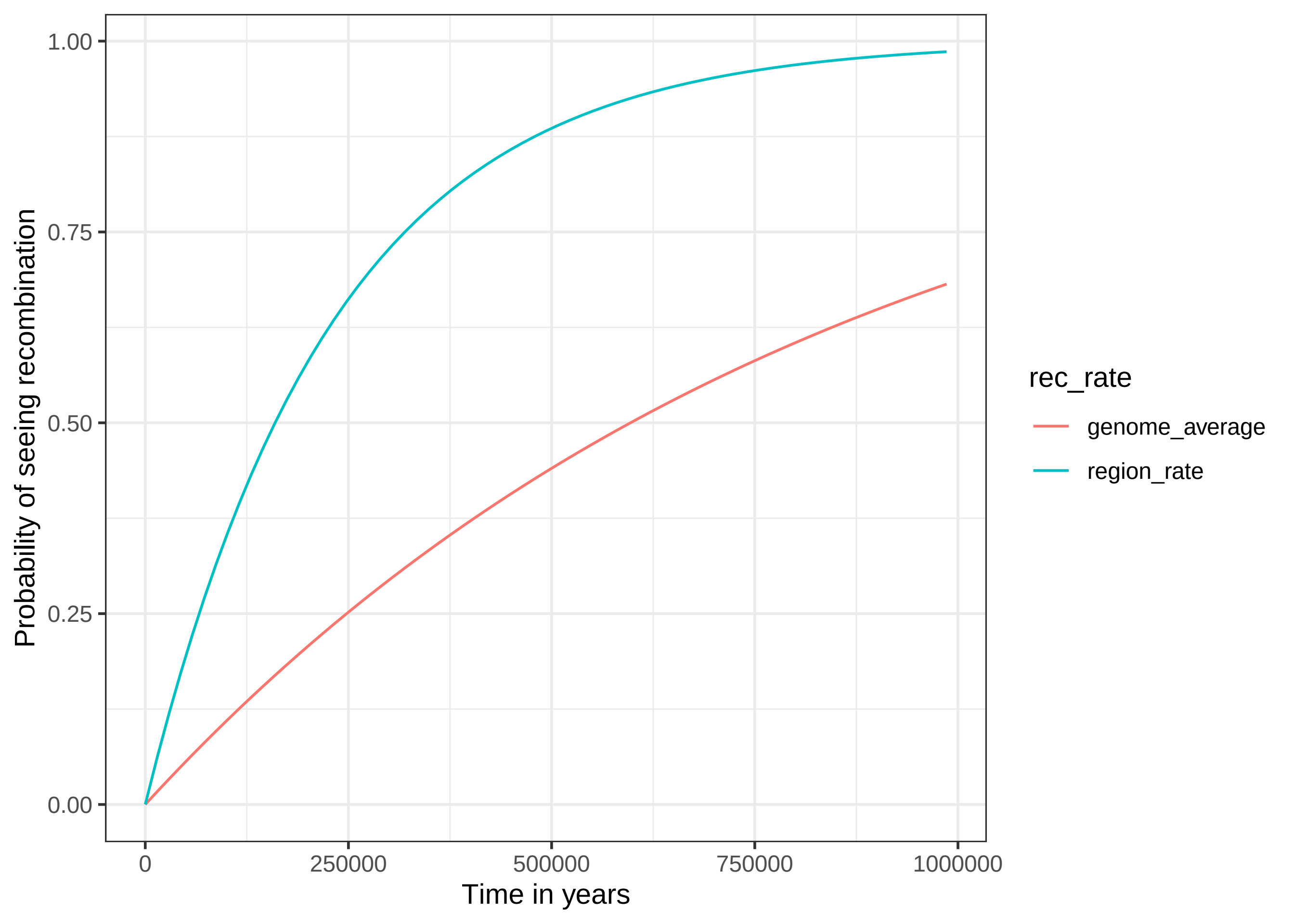

### Fig3_main.pdf

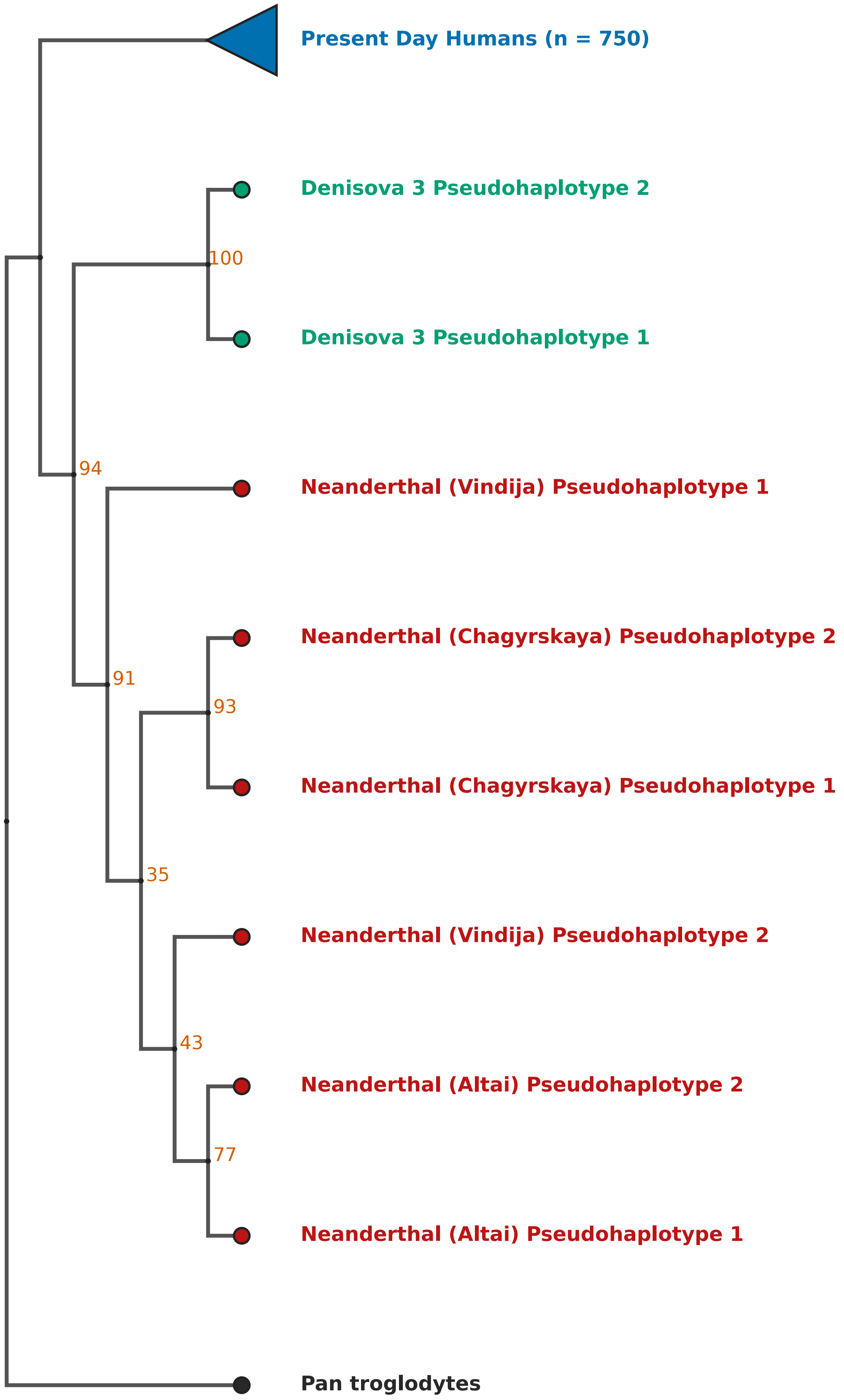
